# The ribosomal protein L22 binds the *MDM4* pre-mRNA and promotes exon skipping to activate p53 upon nucleolar stress

**DOI:** 10.1101/2023.12.29.573614

**Authors:** Jennifer Jansen, Katherine E. Bohnsack, Susanne Böhlken-Fascher, Markus T. Bohnsack, Matthias Dobbelstein

## Abstract

The tumor suppressor p53, along with its antagonists MDM2 and MDM4, represents a central integrator of stress signaling. While DNA damage is the most widely explored trigger of a p53 response, stress arising from dysbalanced assembly of ribosomes in nucleoli is also linked to p53 induction. Deletions of the gene encoding the ribosomal protein L22 (RPL22; eL22) correlate with the presence of full-length *MDM4* mRNA in human cancer, but the mechanistic basis for this phenomenon was hitherto unknown. Here we show that L22, under conditions of ribosomal and nucleolar stress, promotes the skipping of exon 6 within the *MDM4* pre-mRNA. Upon L22 depletion, more full-length MDM4 is maintained, independent of treatment with nucleolar stressors, leading to diminished p53 activity and enhanced cellular proliferation. Mechanistically, L22 binds to specific RNA elements within intron 6 of *MDM4* that correspond to a stem-loop consensus, leading to the skipping of exon 6. This intronic RNA overlaps with the region responsible for splice regulation by ZMAT3. Targeted deletion of these intronic elements largely abolishes L22-mediated exon skipping and re-enables cell proliferation, despite nucleolar stressors such as 5-fluorouracil. L22 also governs alternative splicing of the *L22L1* (*RPL22L1*) and *UBAP2L* mRNAs. Thus, L22 serves as a signaling intermediate that integrates different layers of gene expression. Defects in ribosome synthesis lead to specific alternative splicing, ultimately triggering p53-mediated transcription and arresting cell proliferation.

## INTRODUCTION

p53 is the most frequently mutated tumor suppressor in human cancer (Kandoth et al., 2013; Levine and Oren, 2009; Olivier et al., 2010). When not affected by mutations, the transcription factor p53 is kept in check by its antagonist MDM2, which is itself encoded by a p53-responsive gene (Juven et al., 1993; Leach et al., 1993; Oliner et al., 2016; Wu et al., 1993). MDM2 binds, ubiquitinates and thus destabilizes p53, thereby providing negative feedback (Haupt et al., 1997; Honda et al., 1997; Kubbutat et al., 1997). This system serves to integrate stress signals such as the kinase cascade triggered by DNA damage (Vousden and Prives, 2009). The kinases ATM/ATR and CHK2/CHK1 phosphorylate both p53 and MDM2, thus decreasing MDM2-mediated ubiquitination of p53 and enhancing p53 levels and activity (Li and Kurokawa, 2015; Rizzotto et al., 2021).

Besides DNA damage, nucleolar stress (also known as ribosome assembly stress, triggering the impaired ribosome biogenesis checkpoint) represents an important inducer of p53 (Bursać et al., 2021; Domostegui et al., 2021), but the underlying mechanisms remain to be fully explored. Nucleolar stress occurs when ribosome assembly is impaired, e.g. by alterations in ribosomal RNA (rRNA) synthesis, or by the failure to synthesize or correctly assemble a specific ribosomal component. Under such circumstances, nucleolar proteins, including ribosomal proteins, accumulate in the nucleoplasm. This can induce a p53 response in multiple ways. For example, the 5S ribonucleoprotein complex (5S RNP), consisting of the 5S rRNA and the ribosomal proteins L5 (RPL5; uL18) and L11 (RPL11; uL5), associates with MDM2 and decreases p53 ubiquitination (Boulon et al., 2010; Sloan et al., 2013; Yang et al., 2018).

Besides MDM2, the MDM4 protein is an essential regulator of p53, and the deletion of *MDM4* is embryonic lethal unless accompanied by a deletion of *TP53* (Parant et al., 2001). MDM4 binds both p53 and MDM2, and it supports MDM2-mediated ubiquitination of p53 (Gu et al., 2002; Shvarts et al., 1996; Tanimura et al., 1999). Thus, MDM4 can also serve as an integrator of stress signals to trigger p53. One of the mechanisms that modulate MDM4 in response to stress consists of the skipping of exon 6 in the *MDM4* pre-mRNA. When alterations in splicing lead to elimination of this exon from the pre-mRNA, only a small and unstable form of MDM4 is synthesized, whereas inclusion of exon 6 allows the synthesis of full-length, stable and active MDM4. As a consequence, conditions that increase exon 6 skipping enhance p53 activity (Bardot and Toledo, 2017; Dewaele et al., 2016; Wu et al., 2021).

Massive gene expression analyses in tumors revealed a positive correlation between the skipping of exon 6 in *MDM4* and the presence of two intact alleles encoding the ribosomal protein L22 as one of the most striking correlations (Ghandi et al., 2019). This suggests that L22 and MDM4 might affect each other’s expression or activity. Interestingly, our earlier research revealed that L22 binds to a specific stem-loop motif in RNA, consisting of a G-C base pair on top of the stem, and a U at the 3’ end of the loop. This motif is found in viral RNAs, but its role in mammalian RNA is less well-defined (Dobbelstein and Shenk, 1995). Research in non-mammalian vertebrates suggested a role for L22 in splicing regulation (Zhang et al., 2017). Moreover, low *L22* expression levels correlate with worse outcome of myelodysplastic syndrome in patients and in a mouse model (Harris et al., 2023). Taken together, these observations raise the possibility that the *MDM4* mRNA precursor might contain L22 binding motifs, thus allowing L22 to influence *MDM4* expression.

Here we show that L22 binds to three cognate elements within intron 6 of the *MDM4* pre-mRNA. L22 promotes skipping of exon 6 and thus prevents the synthesis of stable and active, full-length MDM4, thereby increasing the expression of p53 target genes and diminishing cell proliferation. Thus, L22 provides a link between three different aspects of gene expression, i) its extra-ribosomal pool is increased when ribosome biosynthesis is impaired, ii) it then regulates the splicing of *MDM4* and iii) it ultimately triggers the activation of p53 as a transcription factor of specific mRNAs. This signaling mechanism supports cells in responding to nucleolar stress by activating a key tumor suppressive mechanism.

## RESULTS

### RNA polymerase I inhibition triggers L22-dependent *MDM4* exon 6 skipping and reduced L22L1 synthesis

To test the impact of L22 on *MDM4* pre-mRNA splicing, we first induced nucleolar stress in Retinal Pigment Epithelial (RPE) cells, a widely used non-transformed cell line, by treating them with an inhibitor of RNA polymerase I (RNA Pol I), the enzyme that synthesizes three of the four ribosomal RNAs. As expected, depletion of MDM4 induced the expression of the p53 target genes *CDKN1A/p21* and *BBC3/Puma* in these cells (Suppl. Figure 1). In parallel to RNA Pol I inhibition with the small compound BMH-21 (Peltonen et al., 2014), we efficiently depleted L22 by siRNA transfection (Figure 1A, B). As expected (O’Leary et al., 2013), depletion of L22 triggered a compensatory increase in the mRNA level of the L22 paralogue L22L1 (Figure 1A). RNA Pol I inhibition reduced the level of full-length *MDM4* mRNA while increasing expression of the short isoform in which exon 6 is skipped. Strikingly, however, depletion of L22 dramatically diminished this short isoform and restored expression of full-length *MDM4* mRNA, even when nucleolar stress was induced by RNA Pol I inhibition (Figure 1A). Consistent with this, *L22* knockdown also increased the level of MDM4 full-length protein and prolonged its presence despite nucleolar stress (Figure 1B).

**Figure 1:**
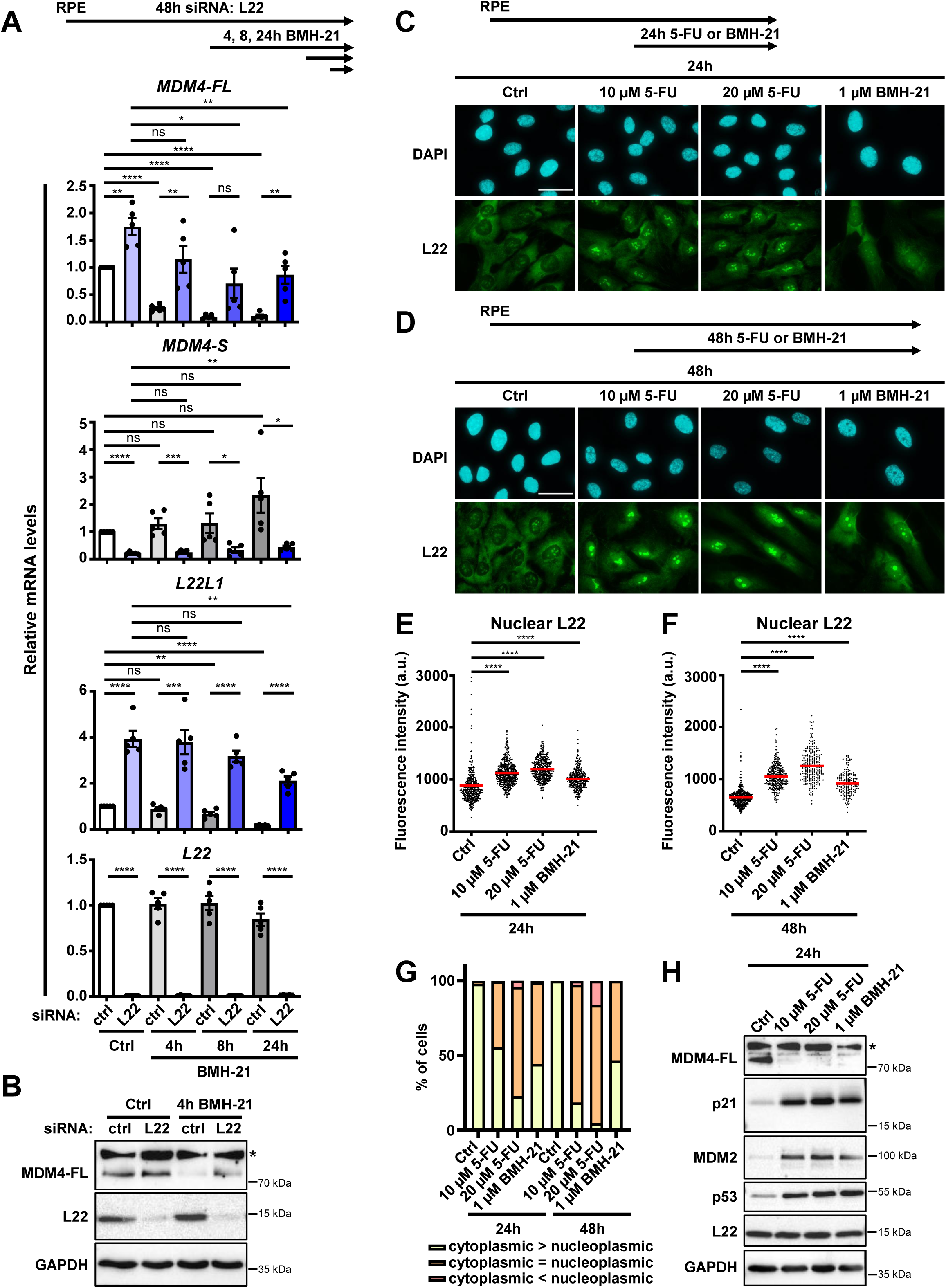
Regulation of *MDM4* and *L22L1* pre-mRNA splicing via L22 upon inhibition of RNA polymerase I. **A.-B.** RPE cells were transfected with siRNA to deplete L22 or with control (ctrl) siRNA for 48 h. Treatment with 1 µM BMH-21 was performed for the indicated periods to inhibit RNA polymerase I and cause nucleolar stress. Ctrl, no BMH-21 treatment. **A.** RT-qPCR analyses using primers against the indicated sequences were performed. Expression levels were normalized relative to the mRNA levels of *36B4* (housekeeping gene) and then to the control sample. The average of five biological replicates is depicted. The statistical significance was assessed by an unpaired t test: ns, not significant; *, P≤0.05; **, P≤0.01; ***, P≤0.001; ****, P≤0.0001. Error bars, standard error of the mean. **B.** Western blot analysis of the indicated proteins was performed, with GAPDH as a sample control. *, background band. **C.-H.** RPE cells were treated with 10 or 20 µM 5-fluorouracil (5-FU) or 1 µM BMH-21 for 24 h or 48 h to induce nucleolar stress. **C.-G.** Immunofluorescence was performed to detect L22. **C.-D.** Representative images (100x objective, bar: 40 µm) of the nuclear (DAPI, blue) and L22 (green) staining upon 24 h (**C**) and 48 h treatment (**D**). **E.-F.** Quantification of nuclear intensities of L22 signal in single cells upon 24 h (**E**) and 48 h (**F**) treatment. Red lines represent the mean fluorescence intensity values. A minimum of 180 cells were quantified for each condition. The statistical significance was assessed by a Mann-Whitney U test. **G.** Quantification of cytoplasmic versus nucleoplasmic intensity of the L22 signal. At least 190 cells were analyzed in each case. **H.** Western blot analysis monitoring the levels of the indicated proteins upon induction of nucleolar stress by treatment with 10 or 20 µM 5-FU or 1 µM BMH-21 for 24 h. *, background band.

Moreover, treatment with BMH-21, but also with a medically relevant inhibitor of rRNA maturation, 5-fluorouracil (5-FU) (Burger et al., 2010), caused redistribution of L22, which is predominantly present in the nucleolus and cytoplasm in untreated cells, to enhance its presence within the nucleoplasm. Specifically, we observed increased total nuclear intensity and a greater nucleoplasmic versus cytoplasmic ratio of the L22 signal upon treatment with BMH-21 or 5-FU (Figures 1C-G). Both BMH-21 and 5-FU reduced the levels of MDM4 full-length protein (Figure 1H). Treatment with these compounds also caused p53 accumulation and augmented the levels of p53 target gene products, p21 and MDM2, while not affecting total L22 protein levels (Figure 1H). Nucleoplasmic enrichment of L22 was also seen upon treatment with BMH-21 or 5-FU in a stably transfected HEK293 Flp-In cell line engineered for the tetracycline-inducible expression of L22 with a C-terminal FLAG tag (Suppl. Figure 2A-C).

We also investigated the impact of L22L1 on expression of *MDM4* isoforms in RPE cells and found that its knockdown induced similar splicing changes as depletion of L22, i.e. decreased expression of the short isoform and increased expression of the full-length mRNA (Suppl. Figure 1). In HEK293T cells, *L22* knockdown again reduced the levels of the short *MDM4* isoform and led to increased full-length *MDM4* mRNA and protein levels (Suppl. Figure 3A, B). In these cells, however, the effect of *L22L1* knockdown on *MDM4* splicing was much less pronounced, suggesting that, in this system, L22 has a stronger role than L22L1 in regulating *MDM4* pre-mRNA splicing. To further consolidate the influence of L22 and L22L1 on *MDM4* isoform expression, the paralogous ribosomal proteins were overexpressed in HEK293T cells, which are particularly suited for plasmid transfection. Reciprocally to the effects of the knockdowns, this revealed enhanced *MDM4* exon skipping, indicated by reduced expression of the full-length *MDM4* mRNA and increased levels of the short *MDM4* isoform (Suppl. Figure 3C, D). Taken together, these results strongly suggest that L22 modulates *MDM4* exon skipping in the context of nucleolar stress.

### Depletion of the ribosomal protein S13 triggers L22-dependent nucleolar stress, *MDM4* exon skipping, and p53 activation

Next, we induced nucleolar stress independent of drugs, by depletion of a ribosomal protein that belongs to the small ribosomal subunit, S13 (RPS13; uS15), alone or together with L22 depletion (Figure 2A-B). The depletion of S13 alone led to L22 accumulation in the nucleoplasm (Figure 2C-E and Suppl. Figure 4A-C). Lack of S13 alone also diminished full-length *MDM4* mRNA, increased the levels of the short *MDM4* isoform, and triggered the expression of the p53-responsive genes *p21*, *MDM2* and *Puma* (Figure 2A). However, when L22 was depleted along with S13, these effects were substantially diminished (Figure 2A). L22 co-depletion also restored the presence of detectable MDM4 protein upon *S13* knockdown, and it reduced the protein levels of the p53-responsive gene products p21 and MDM2 (Figure 2B).

**Figure 2:**
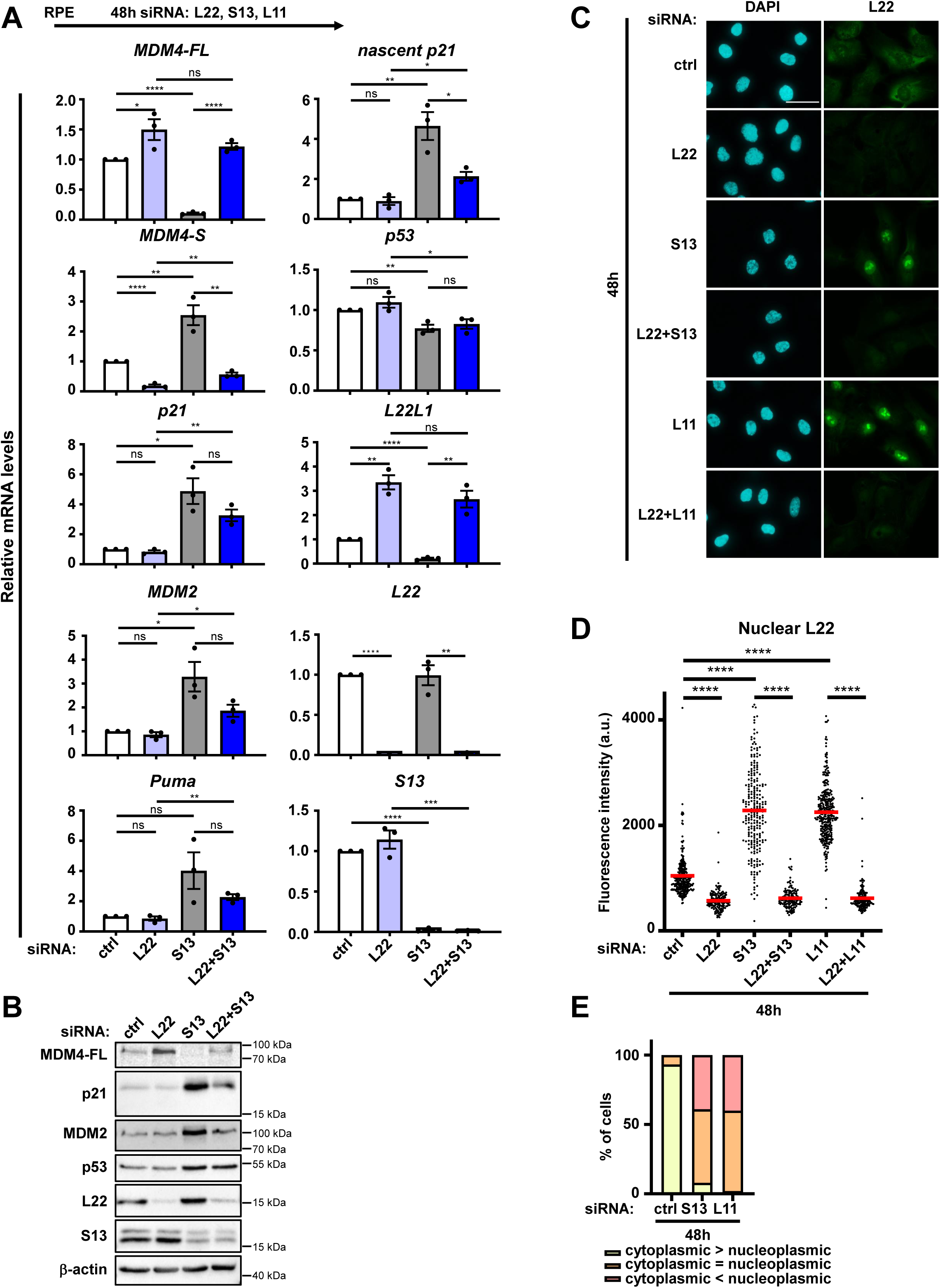
p53 activity upon nucleolar stress, dependent on L22. **A.-B.** RPE cells were transfected with siRNA to deplete L22 and/or S13 (the latter to induce nucleolar stress), for 48 h. **A.** RT-qPCR analyses of the indicated mRNAs were performed. Expression levels were normalized to the mRNA levels of *36B4*. Statistical significance: ns, not significant; *, P≤0.05; **, P≤0.01; ***, P≤0.001; ****, P≤0.0001. **B.** Western blot analysis of the proteins indicated was performed, using β-actin as a sample control. **C.-E.** RPE cells were transfected with siRNA to deplete L22 and/or S13 or L11 for 48 h. Immunofluorescence staining of L22 was performed. **C.** Representative images (100x objective, bar: 40 µm) of the nuclear (DAPI, blue) and L22 (green) staining. **D.** Quantification of single nuclear intensities of L22 signal. Red lines represent the mean fluorescence intensity values. A minimum of 130 cells were quantified for each condition. **E.** Quantification of cytoplasmic versus nucleoplasmic intensity of L22 signal. At least 150 cells were analyzed for each condition.

We also induced nucleolar stress by depleting the ribosomal proteins L5 and L11, which are part of the large ribosomal subunit sub-complex, the 5S RNP. As the 5S RNP binds and inhibits MDM2, this raises the question whether the depletion of these ribosomal proteins decreases p53 activity through free MDM2, or whether this still activates p53 through removal of full-length MDM4. To test this, we combined the depletion of L5 or L11 with *L22* knockdown (Suppl. Figure 4D, E). Notably, *L5* knockdown caused a reduction of the L11 protein level and vice versa, presumably due to mutual stabilization of the two proteins within the 5S RNP (Suppl. Figure 4E). Depleting the 5S RNP led to the nucleoplasmic accumulation of L22 (Figure 2C-E and Suppl. Figure 4A-C). Similar to the depletion of S13, depleting L5 or L11 reduced the levels of full-length *MDM4* mRNA and protein, and it increased the expression of p53 target genes (Suppl. Figure 4D, E). Finally, removing S13 or L11 caused a similar L22 re-distribution in a stably transfected HEK293 Flp-In cell line expressing FLAG-tagged L22 (Suppl. Figure 5A-C). In conclusion, L22 not only mediates exon-skipping of *MDM4* but also contributes to the subsequent activation of p53 upon depletion of other ribosomal proteins.

### Intron 6 of the *MDM4* pre-mRNA contains three consensus L22 binding sites and their deletion abolishes the impact of L22 on *MDM4* splicing

L22 has long been known as a ligand of the Epstein Barr Virus Expressed RNAs (EBERs) (Toczyski et al., 1994; Toczyski and Steitz, 1991), and in vitro selection has revealed a specific stem-loop RNA motif that binds L22 (Dobbelstein and Shenk, 1995) (Figure 3A). As L22 induces *MDM4* exon skipping, a direct interaction of L22 with the pre-mRNA of *MDM4*, perhaps through similar RNA motifs, is possible. Consistent with this hypothesis, subjecting the RNA sequence around exon 6 to a folding algorithm, i.e. the RNAfold web server minimum free energy (MFE) function (http://rna.tbi.univie.ac.at/cgi-bin/RNAWebSuite/RNAfold.cgi), revealed the presence of three L22-binding consensus motifs within intron 6, in close proximity to each other and to the adjacent exon 6 (Figure 3B, C). Strikingly, all of the three sequences are conserved in the *MDM4* homologous gene within the murine genome (Suppl. Figure 6A). We therefore tested whether the portion of intron 6 that contains the L22-binding motifs is required for L22-mediated alternative splicing. To this end, we removed this portion of the intron in RPE cells by CRISPR/Cas9 genome editing (Figure 3B, red scissors; Suppl. Figure 6B). Cells that carry the deletion in a homozygous fashion remained viable. When these cells were subjected to nucleolar stress via S13 depletion, *MDM4* pre-mRNA splicing was not affected, and L22 depletion did not detectably change the splice pattern of *MDM4* either (Figure 3D-G). In line with this, cells lacking the three L22 consensus sequences showed reduced p53 activity upon S13 depletion, as evidenced by diminished expression of *p21*, *MDM2* and *Puma* (Suppl. Figure 6C, D). This demonstrates that the region of the *MDM4* pre-mRNA containing the consensus L22 binding sites is strictly required for the impact of L22 on *MDM4* splicing.

**Figure 3:**
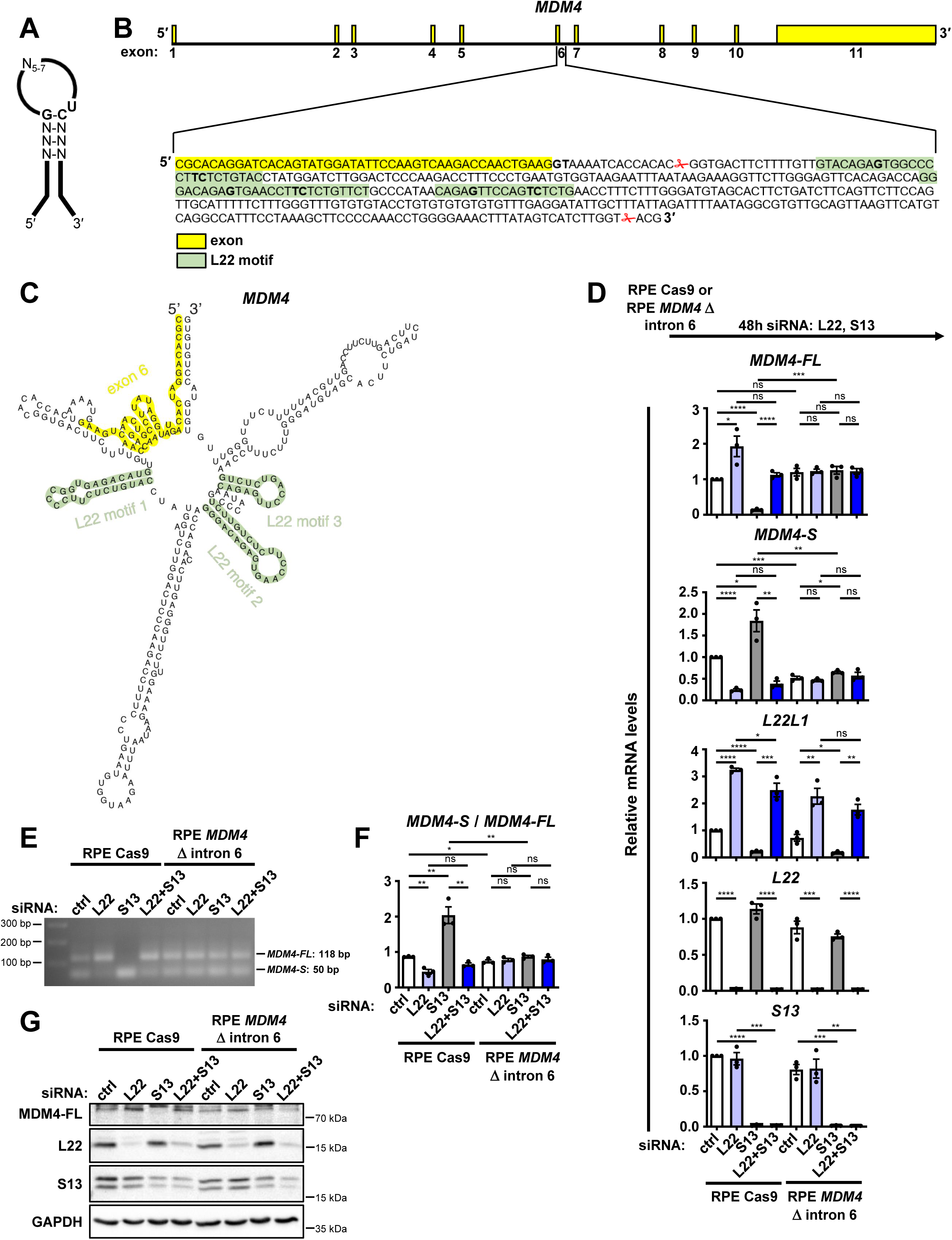
Abolished splice regulation by L22 upon deletion of L22-binding consensus sequences from *MDM4* intron 6. **A.** RNA motif that mediates the interaction with L22: a stem-loop structure with a G–C base pair (bp) at the top of the stem (G on the 5’ side, C on the 3’ side) and U at the 3’ end of the loop. The conserved bases G, C and U are marked in bold. Adapted from Dobbelstein and Shenk, 1995. **B.** Schematic view of the *MDM4* gene with correct proportions of exons and introns, with a magnified view of the last 46 bp of exon 6 (marked in yellow) and the first 387 bp of intron 6. Sequences in intron 6 predicted to form L22-binding stem loops are marked in green, and the characteristic bases G, C and U (T) are indicated in bold. Red scissor symbols show cleavage sites for CRISPR/Cas9-mediated deletion of all three L22-binding stem loop sequences in *MDM4* intron 6. **C.** RNA folding prediction of the last 46 bp of exon 6 and the first 268 bp of intron 6 of the *MDM4* gene (i.e., most of the sequence shown in **B**) using the RNAfold web server (http://rna.tbi.univie.ac.at/cgi-bin/RNAWebSuite/RNAfold.cgi) minimum free energy (MFE) function. Exon 6 and sequences predicted to form L22-binding stem loops are marked as in **B**. **D.-G.** RPE Cas9 or RPE *MDM4* Δ intron 6 cells were transfected with siRNA to deplete L22 and/or S13 for 48 h. **D.** RT-qPCR analyses using primers against the indicated sequences were performed. Expression levels were normalized to the mRNA levels of *36B4*. Statistical significance: ns, not significant; *, P≤0.05; **, P≤0.01; ***, P≤0.001; ****, P≤0.0001. **E.-F.** RT-qPCR analysis using primers leading to a longer product for *MDM4-FL* and a shorter product for *MDM4-S* was performed. **E.** RT-qPCR products were visualized by agarose gel electrophoresis and staining with SERVA DNA Stain Clear G. **F.** Based on band quantification, *MDM4-S*/*MDM4-FL* ratios were calculated. The average of three biological replicates is depicted. The statistical significance was assessed by an unpaired t test. **G.** Western blot analysis was performed to indicate MDM4 protein levels largely corresponding to the mRNA.

### L22 also regulates the expression of *UBAP2L* and *L22L1* isoforms

The previous analysis of exon inclusion vs. genotype in tumor cells not only revealed a correlation between *L22* deletions and full-length *MDM4* expression, but also between *L22* deletions and expression of the gene *UBAP2L* (Ghandi et al., 2019), which encodes a strongly conserved ubiquitin– and RNA-binding protein (Guerber et al., 2022). We therefore hypothesized that L22 might also act as a regulator of *UBAP2L* splicing, and we investigated whether this is the case by combining L22 and S13 depletions (Figure 4A-C). Here, L22 depletion strongly reduced the abundance of a transcript isoform (UBAP2L-201) that contains the last *UBAP2L* exon, exon 27, in a more distal position (Figure 4A, B). Depleting L22 rather enhanced the inclusion of a more proximal exon 27, corresponding to the isoform UBAP2L-207 that is considered the major one (Figure 4A, B), thereby enhancing the protein levels of this main UBAP2L version (Figure 4C). This might be explained by the presence of L22 binding sites within the above-mentioned exons (Figure 4B).

**Figure 4:**
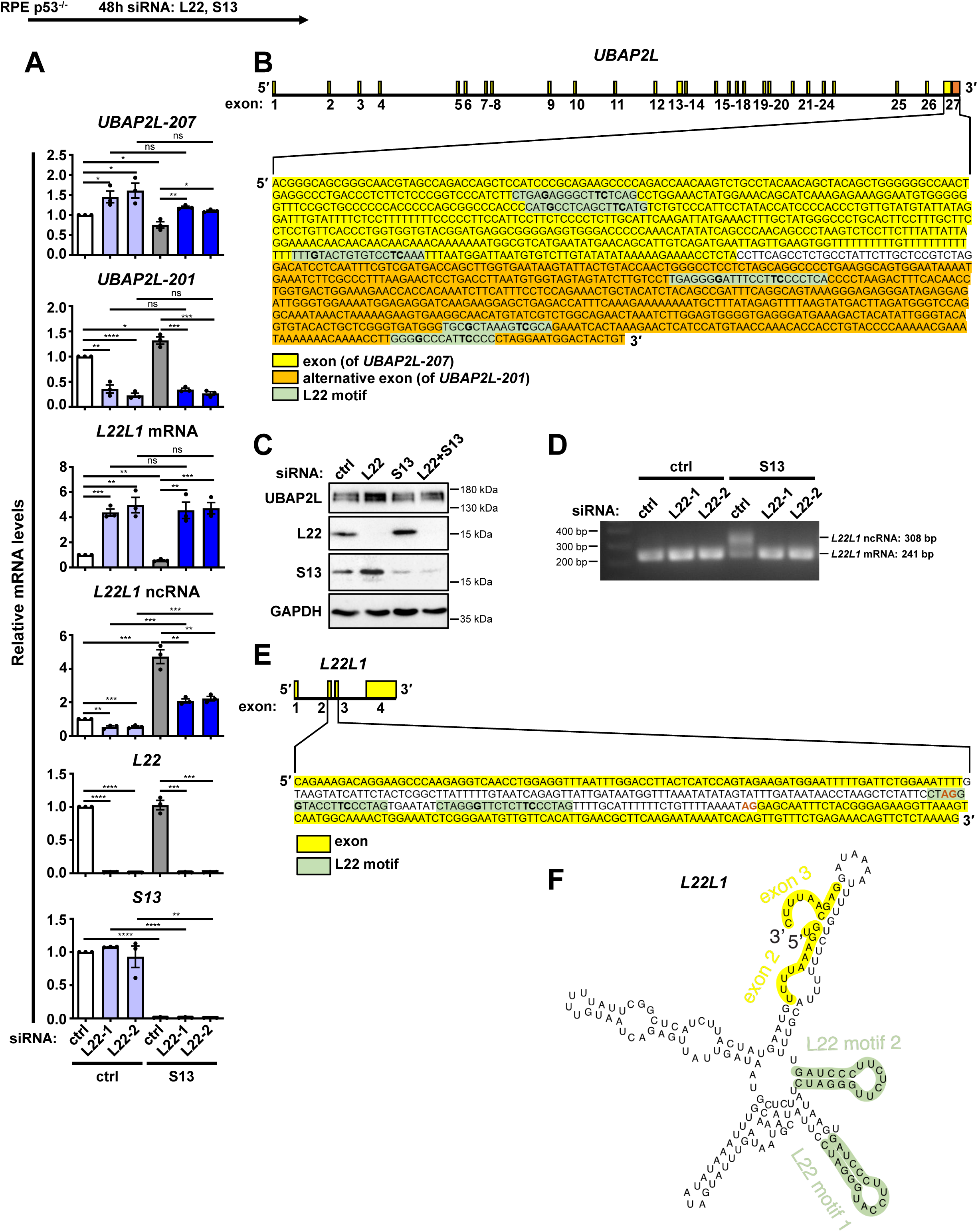
Splice regulation of *L22L1* and *UBAP2L* by L22. **A.** RPE p53^-/-^ cells were transfected with siRNA to deplete L22 and/or S13 as indicated for 48 h. RT-qPCR analyses using primers against the indicated sequences were performed. Expression levels were normalized to the mRNA levels of *36B4*. Statistical significance: ns, not significant; *, P≤0.05; **, P≤0.01; ***, P≤0.001; ****, P≤0.0001. **B.** Schematic view of the *UBAP2L* gene and magnified view of the end of its sequence. The last exon, exon 27, is marked in yellow for the main *UBAP2L* variant *UBAP2L-207*, and in orange for the *UBAP2L* variant *UBAP2L-201*. Within both exons, sequences predicted to form L22-binding stem loops are marked in green, and the characteristic bases G, C and U (T) are indicated in bold. **C.** RPE p53^-/-^ cells were transfected with siRNA to deplete L22 and/or S13 for 48 h. Western blot analysis revealed L22-dependency of UBAP2L protein levels. **D.** RT-qPCR analysis of the same samples as in **A** was performed, using primers leading to a longer product for *L22L1* ncRNA and a shorter product for *L22L1* mRNA. RT-qPCR products were visualized by agarose gel electrophoresis. **E.** Schematic view of the *L22L1* gene. Magnified view: sequence of exons 2 and 3 (marked in yellow) and intron 2. Sequences in intron 2 predicted to form L22-binding stem loops are marked in green as in **B**. The splice acceptor site of intron 2 and an alternative splice acceptor site leading to the formation of the *L22L1* non-coding RNA (ncRNA) variant are indicated in bold and orange. **F.** RNA folding prediction of the last 10 bp of exon 2, intron 2, and the first 10 bp of exon 3 of the *L22L1* gene (i.e., part of the sequence shown in **E**) using the RNAfold web server (http://rna.tbi.univie.ac.at/cgi-bin/RNAWebSuite/RNAfold.cgi) minimum free energy (MFE) function. Exonic sequences and L22-binding motifs are indicated according to the scheme used in **B** and **E**.

We (Figures 1A, 2A, 3D) and others (O’Leary et al., 2013) have observed that L22 depletion increases the levels of *L22L1* mRNA. This raised the question whether L22 also induces alternative splicing of *L22L1*. Indeed, the depletion of L22 enhanced the abundance of a functional mRNA encoding L22L1 and reduced a non-coding variant of this mRNA, while S13 depletion led to increased levels of *L22L1* non-coding RNA (Figure 4A, D). We found juxtaposed L22-binding consensus sites in this case as well (Figure 4E, F). In conclusion, L22 regulates the splicing of at least two more genes.

### ZMAT3 requires an overlapping portion of intron 6 to induce *MDM4* exon skipping, but does not affect the splicing of *L22L1* and *UBAP2L*

In previous studies, the protein ZMAT3 was characterized as a splice factor that also triggers *MDM4* exon skipping (Bieging-Rolett et al., 2020). We therefore asked whether ZMAT3 might act on similar RNA sequence elements as L22 to modulate splicing. To investigate this, we combined L22 and ZMAT3 depletions (Figure 5A-C). Indeed, the depletion of ZMAT3 enhanced the abundance of full-length *MDM4* and reduced the amount of short isoform mRNA. Moreover, the partial deletion of intron 6 (including the consensus L22 binding sites) abolished the response of *MDM4* splicing to ZMAT3, at first glance suggesting a similar splice-regulatory mode for ZMAT3 and L22 (Figure 5A, B). However, the simultaneous depletion of ZMAT3 and L22 diminished the short *MDM4* isoform even more strongly than the single knockdowns, suggesting that they may not only be part of the same splice-regulatory mechanism (Figure 5A, B). Moreover, ZMAT3 depletion did not affect the ratio of *L22L1* and *UBAP2L* isoforms, again suggesting that the activity of L22 and ZMAT3 is not the same (Figure 5A). We propose that ZMAT3 and L22 act through similar portions of the *MDM4* intron 6, but that they still use different and at least partially independent mechanisms for doing so.

**Figure 5:**
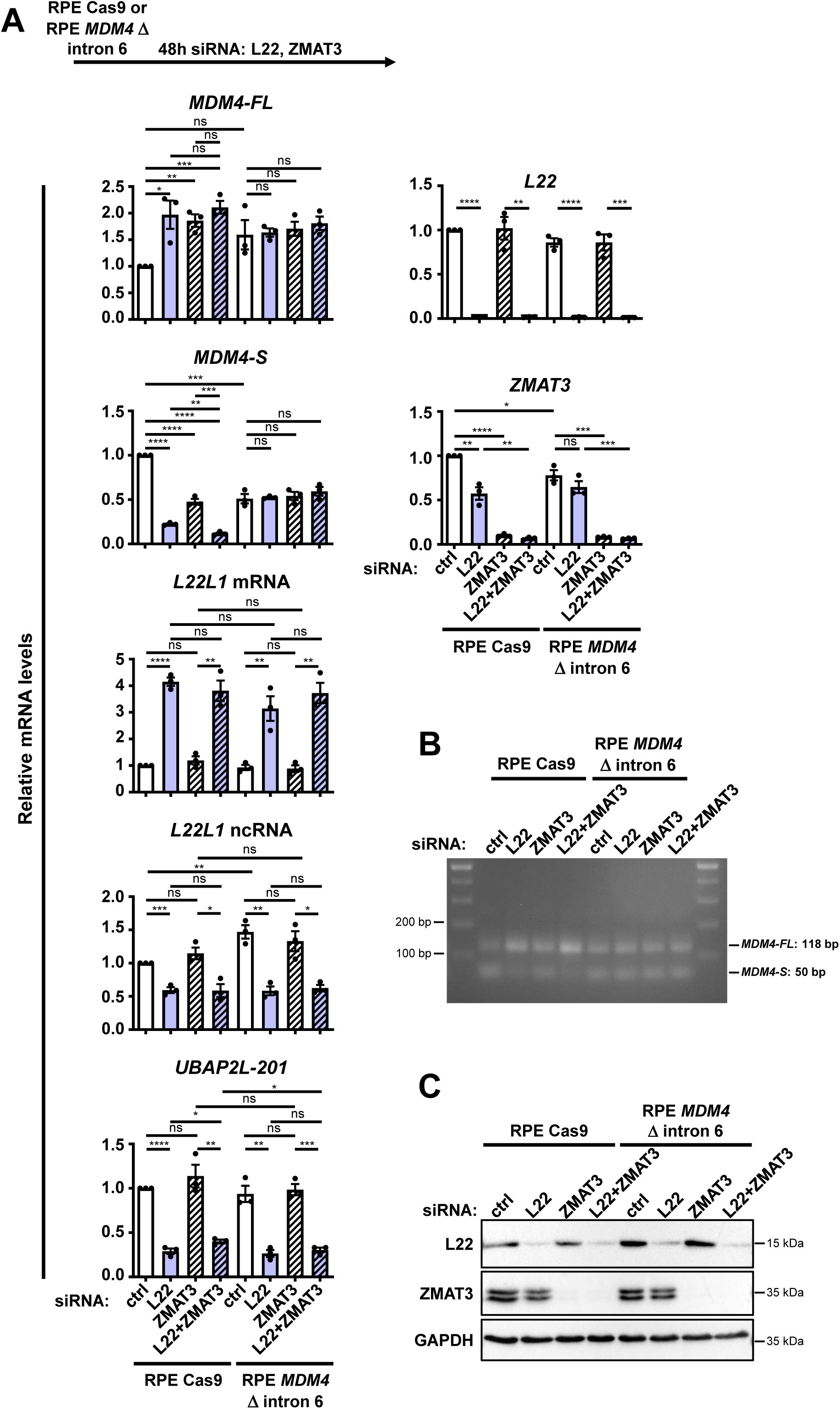
Overlapping intronic sequences required for splice regulation of *MDM4* by L22 and ZMAT3. **A.-C.** RPE Cas9 or RPE *MDM4* Δ intron 6 cells were transfected with siRNA to deplete L22 and/or ZMAT3 for 48 h. **A.** RT-qPCR analyses using primers against the indicated sequences were performed. Expression levels were normalized to the mRNA levels of *36B4*. Statistical significance: ns, not significant; *, P≤0.05; **, P≤0.01; ***, P≤0.001; ****, P≤0.0001. **B.** RT-qPCR analysis using primers leading to a longer product for *MDM4-FL* and a shorter product for *MDM4-S* was performed. RT-qPCR products were visualized by agarose gel electrophoresis. **C.** Western blot analysis of the proteins indicated was performed, using GAPDH as a sample control.

### L22-binding motifs 1 and 2 are especially important for interaction with the *MDM4* pre– mRNA and its splicing

To define more precisely how L22 influences *MDM4* pre-mRNA splicing, we performed reporter assays. The reporter expresses the enhanced green fluorescent protein (eGFP) upon exon skipping. *MDM4* exon 6, along with the flanking intronic regions, were cloned into this reporter (Figure 6A). As expected, siRNA-mediated depletion of L22 led to enhanced exon inclusion and thus diminished eGFP levels (Figure 6B, C and Suppl. Figure 7A). Next, we mutated the three L22 binding motifs of intron 6 within this reporter construct, individually or in combination, or deleted all three motifs. Upon removal of the first and/or second L22 consensus motif, exon skipping became less dependent on, or even entirely independent of, L22 (Figure 6B, C and Suppl. Figure 7A). Hence, these two RNA motifs are of particular importance for pre-mRNA splicing regulation by L22.

**Figure 6:**
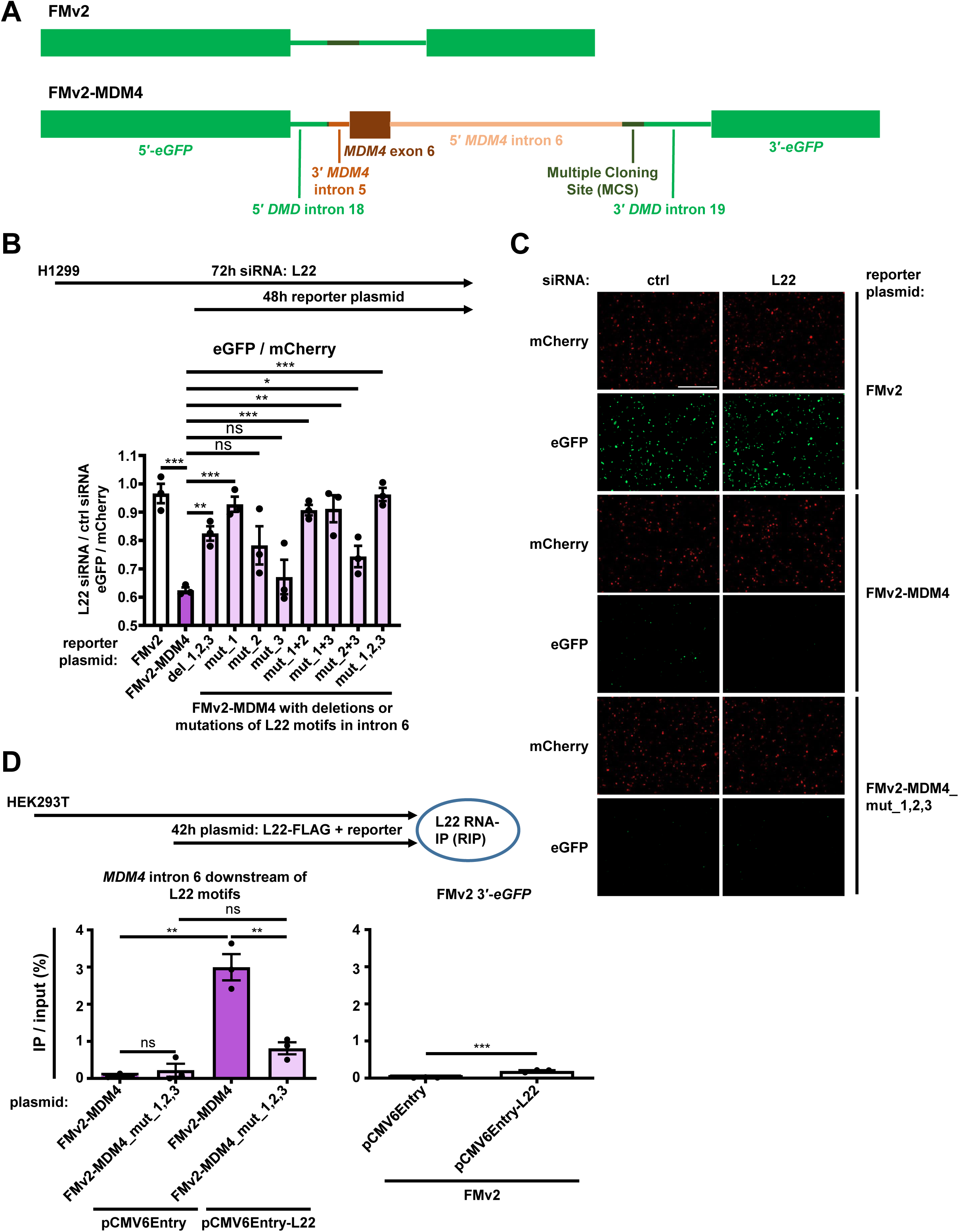
L22 binding to consensus sequences in intron 6 of *MDM4* pre-mRNA. **A.** A splicing reporter assay based on Fukushima et al., 2020 was used to investigate the sequence requirements needed for L22 to bind *MDM4* pre-mRNA and regulate its splicing. To generate the control plasmid FMv2, the following sequences were inserted into the pcDNA3.1/Hygro(+) vector containing the CMV promoter by using the restriction enzymes *Nhe*I and *Xho*I: the mCherry coding sequence, the 5′ end of the *eGFP* gene (5′-*eGFP*), the 5′ end of *DMD* intron 18 (5′ *DMD* intron 18), a Multiple Cloning Site (MCS), the 3′ end of *DMD* intron 19 (3′ *DMD* intron 19), and the 3′ end of the *eGFP* gene (3′-*eGFP*). To generate the *MDM4* splicing reporter plasmid FMv2-MDM4, the following sequences were inserted into the MCS of FMv2 by using the restriction enzymes *Hind*III and *Bam*HI: the 3′ end (34 bp) of *MDM4* intron 5 (3′ *MDM4* intron 5), *MDM4* exon 6 (68 bp), and the 5′ end (398 bp) of *MDM4* intron 6 (5′ *MDM4* intron 6) containing all three previously identified L22 consensus sequences. Upon transfection of either FMv2 or FMv2-MDM4, mCherry is constitutively expressed and serves as a transfection marker. eGFP is constitutively expressed upon FMv2 overexpression, but only if *MDM4* exon 6 is skipped in case of FMv2-MDM4 overexpression, therefore serving as a splicing marker. **B.-C.** FMv2-MDM4 versions with either deletion of all three L22 consensus sequences in *MDM4* intron 6 (FMv2-MDM4_del_1,2,3) or with mutations of the three conserved nucleotides of the single or combined L22 consensus sequences as indicated (FMv2-MDM4_mut_[…]) were generated by site-directed mutagenesis. H1299 cells were transfected with siRNA to deplete L22 for 72 h, and with FMv2, FMv2-MDM4 and the different mutated versions of FMv2-MDM4, at 24 hours after the siRNA transfection, followed by incubation for 48 h. Red fluorescence (mCherry) and green fluorescence (eGFP) were detected. **B.** The eGFP fluorescence intensity among cells with high mCherry intensity (>20 cells per sample) was quantified, and the change of the eGFP/mCherry ratio with L22 compared to control depletion was visualized. The average of three biological replicates is depicted. The statistical significance was assessed by an unpaired t test: ns, not significant; *, P≤0.05; **, P≤0.01; ***, P≤0.001. **C.** Images (5x objective, bar: 800 µm) of the eGFP (green) and mCherry (red) staining. **D.** HEK293T cells were transfected with pCMV6Entry or pCMV6Entry-L22 and with either FMv2, FMv2-MDM4 or FMv2-MDM4_mut_1,2,3 for 42 h, followed by RNA immunoprecipitation (RIP) using FLAG beads to precipitate tagged L22. RT-qPCR analyses using primers against the indicated sequences were performed for the immunoprecipitation (IP) and 5% input samples shown, and IP signal was normalized to input signal.

Next, we addressed whether L22 directly binds to its consensus motifs in intron 6 of the *MDM4* pre-mRNA to promote exon skipping. HEK293T cells were transfected with the wildtype *MDM4* reporter plasmid (containing *MDM4* exon 6 and flanking regions of intron 6), or with constructs that carried mutations of the three L22 motifs, along with an expression construct for FLAG-tagged L22. Subsequently, we performed RNA immunoprecipitation (RIP) using anti-FLAG antibody coupled to beads (Figure 6D and Suppl. Figure 7B, C). *MDM4* RNA corresponding to the region directly downstream of the L22 consensus sequences in intron 6 was co-precipitated with L22 (Figure 6D), and this precipitation was reduced by two thirds if the exogenous *MDM4* RNA contained mutated L22 motifs (Figure 6D). Thus, L22 binds to its consensus motifs in the *MDM4* pre-mRNA to regulate *MDM4* isoform expression.

### Removing L22-binding sites within intron 6 of the *MDM4* pre-mRNA suppresses p53 activation and enables cell proliferation during nucleolar stress

Finally, we sought to clarify how L22-mediated splicing regulation of the *MDM4* pre-mRNA influences p53 activity and cell proliferation, especially when cells are confronted with a clinically relevant compound inducing nucleolar stress, 5-FU (Burger et al., 2010). We treated RPE cells, with or without the critical region within *MDM4* intron 6 containing the L22 binding motifs, with 5-FU, followed by quantification of cell proliferation (Figure 7A-D), and monitoring of p53-responsive gene expression (Figure 7E). As expected, 5-FU induced a p53 response, and it compromised cell proliferation during the following days. However, when the L22-binding sites within *MDM4* intron 6 had been removed, the cells not only displayed reduced expression of p53-responsive genes, but they also proliferated to a significantly higher extent than the parental cells (Figure 7A-E). As expected, 5-FU induced *MDM4* exon 6 skipping and reduced full-length *MDM4* mRNA expression only if the L22 consensus sequences in *MDM4* intron 6 were present (Figure 7E-G). By contrast, the reduction of *L22L1* expression upon 5-FU treatment was similar in both cell lines (Figure 7E). In conclusion, L22 serves to integrate nucleolar stress signaling into growth arrest, reaching from compromised rRNA synthesis or processing, through alternative *MDM4* splicing and p53-mediated gene expression, to a pronounced block in cell proliferation (Figure 8).

**Figure 7:**
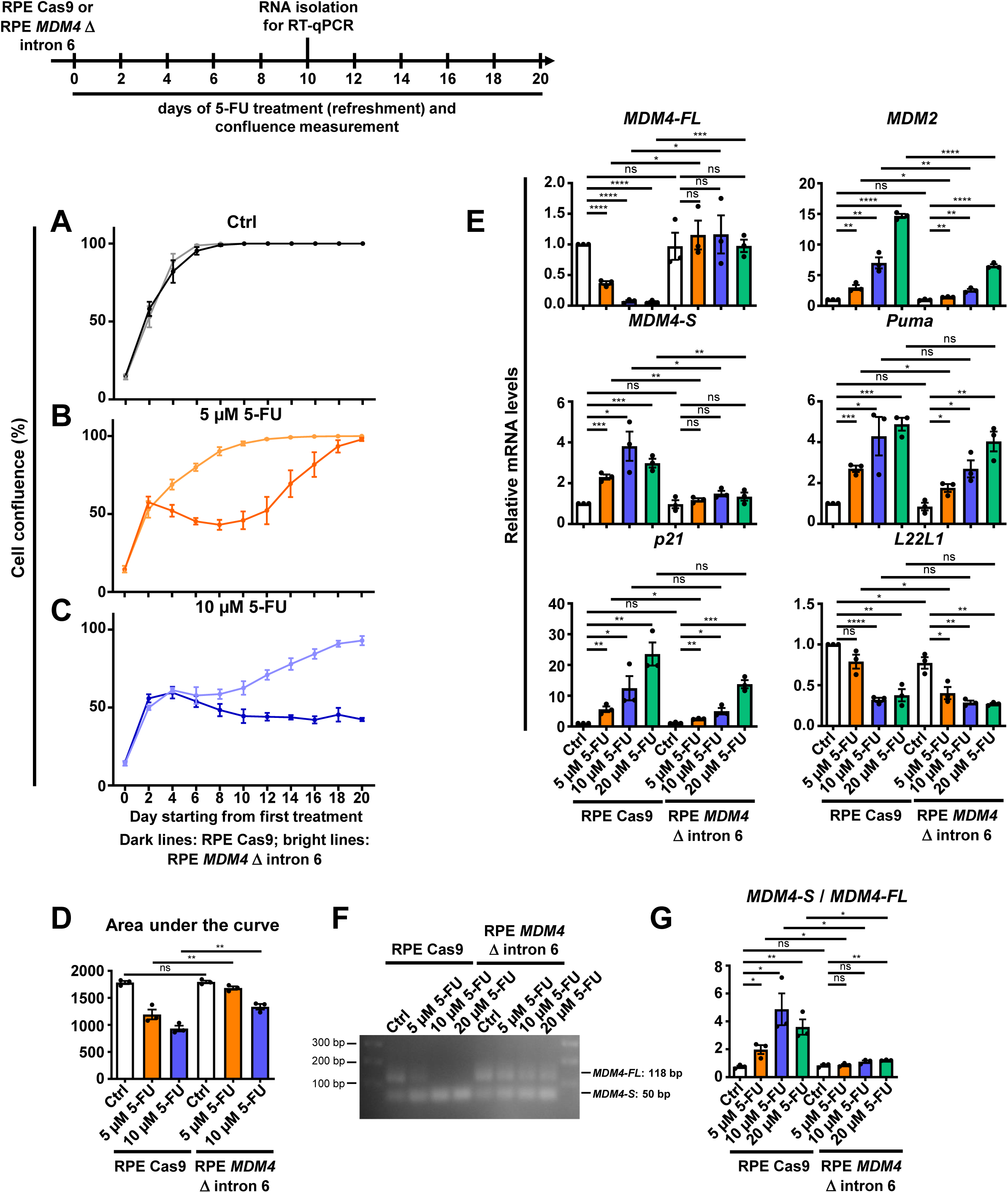
Resistance towards 5-fluorouracil upon abrogation of *MDM4* splicing regulation by L22. **A.-D.** RPE Cas9 (dark) or RPE *MDM4* Δ intron 6 (light) cells were treated with 5-fluorouracil (5-FU) or control-treated (ctrl) as indicated for 20 days. Every other day, the treatment was refreshed and cell confluence was measured. **A.-C.** For each condition, results of three biological replicates (with three technical replicates each) are depicted. **D.** Statistical analysis was performed by comparing the area under the curve (AUC) values, which were calculated from the confluence graphs (**A-C**), of both cell lines using an unpaired t test: ns, not significant; *, P≤0.05; **, P≤0.01; ***, P≤0.001; ****, P≤0.0001. **E.-G.** RPE Cas9 or RPE *MDM4* Δ intron 6 cells were treated with 5-FU as in **A-D**. After 10 days of treatment, cells were harvested for RNA isolation. **E.** RT-qPCR analyses to quantify mRNA levels corresponding to the indicated genes were performed, normalized to *36B4*. **F.-G.** RT-qPCR analysis using primers leading to a longer product for *MDM4-FL* and a shorter product for *MDM4-S* was performed. **F.** RT-qPCR products were visualized by agarose gel electrophoresis. **G.** Based on band quantification, the *MDM4-S*/*MDM4-FL* ratio was calculated.

**Figure 8:**
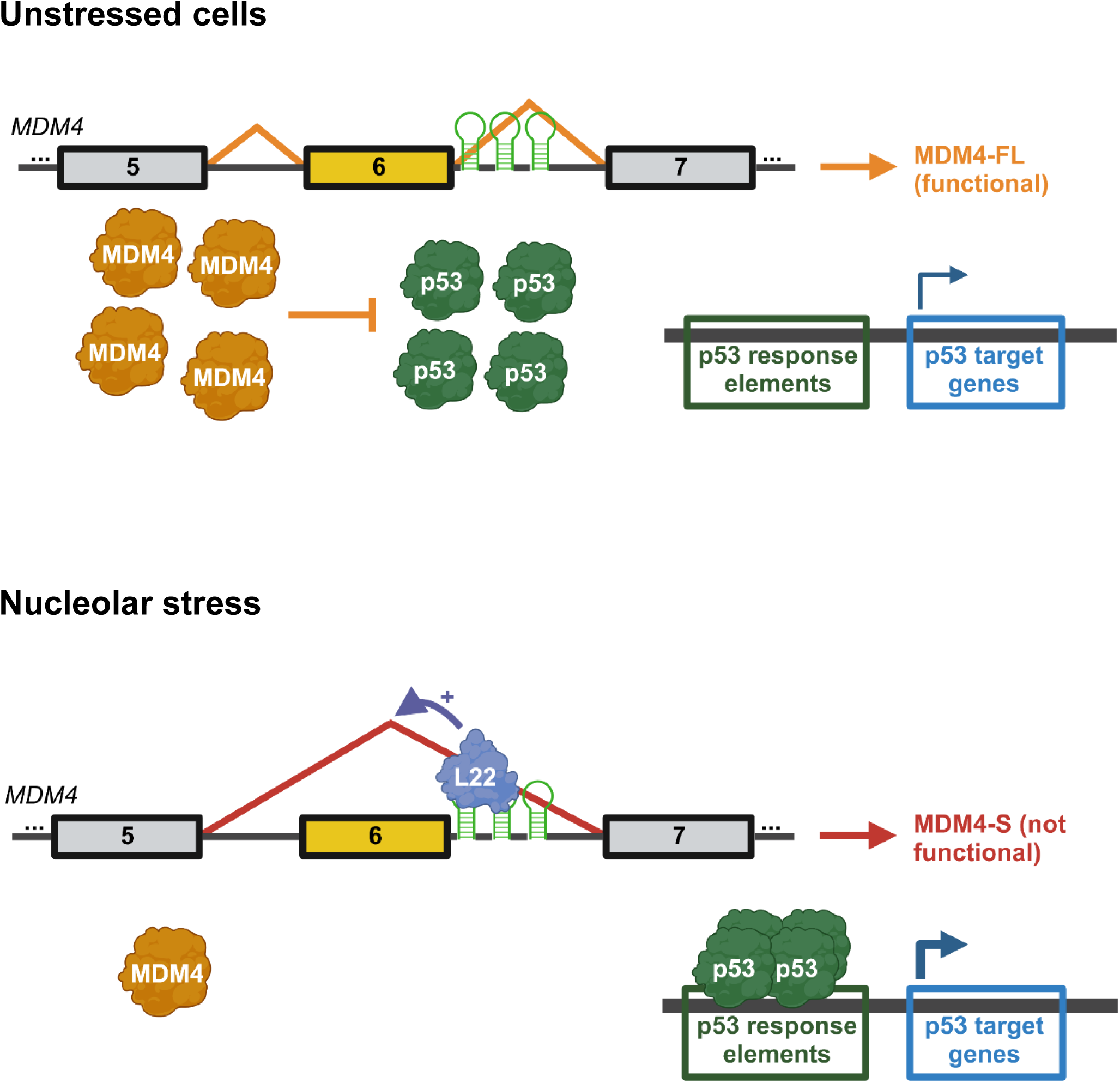
Alternative *MDM4* splicing and p53 activation through L22 upon nucleolar stress. L22, by promoting alternative splicing of *MDM4* pre-mRNA, activates p53-mediated gene expression and thus translates nucleolar stress into growth arrest.

## DISCUSSION

Our results identify L22 as a signaling transmitter that triggers a p53 response upon impaired ribosome synthesis. When biogenesis of the translation machinery is perturbed, L22 accumulates in the nucleoplasm and associates with a cognate binding motif within a subset of pre-mRNAs, including that of *MDM4*. Splicing of the *MDM4* pre-mRNA is adjusted towards the skipping of exon 6, leading to increased expression of a truncated and poorly functional isoform of *MDM4*. This promotes p53 to activate the transcription of its target genes.

Interestingly, a link between L22 and alternative splicing of the *MDM4* pre-mRNA was recently reported in the context of colon cancer of the microsatellite instability subtype (Weinstein et al., 2023), implying that our findings in RPE and HEK293T cells are broadly applicable. These complementary studies both also reveal a role for L22 in modulating the splicing of the *L22L1* pre-mRNA. Importantly, our work provides additional mechanistic details on the impact of nucleolar stress in this context, on the critical nucleotides within the L22-binding motifs in the *MDM4* pre-mRNA, and on the role of ZMAT3. Moreover, we identify the *UBAP2L* pre-mRNA as an additional target of L22-modulated alternative splicing.

Nucleolar stress is a widely encountered cell condition. It is not only triggered by dysbalanced ribosomal protein synthesis, e.g. in aneuploid cells, but also occurs in response to DNA damage, as the DNA encoding rRNA responds to damage by perturbed transcription and disintegration of nucleoli (Kruhlak et al., 2007; Rubbi and Milner, 2003). A number of chemotherapeutics can trigger nucleolar stress, including 5-FU used in our study, but also oxaliplatin (Burger et al., 2010; Hein et al., 2013). 5-FU causes nucleolar stress through incorporation into rRNA (Ghoshal and Jacob, 1997; Sun et al., 2007). Oxaliplatin differs from cisplatin by its ability to modify rRNA, whereas cisplatin primarily acts on DNA (Bruno et al., 2017; Sutton and DeRose, 2021). These agents are thus plausible triggers of L22-mediated *MDM4* exon skipping.

Alongside L22, the p53-inducible factor ZMAT3, the arginine methyltransferases PRMT5 and PRMT1, and the splicing factor PRPF19 can modulate the splicing of *MDM4* (Bezzi et al., 2013; Bieging-Rolett et al., 2020; Jackson-Weaver et al., 2020; Yano et al., 2021). This emphasizes the role of *MDM4* exon skipping in integrating the responses to multiple stresses, all leading to p53 activation. Our data suggest, however, that L22 is the first factor found to transmit nucleolar stress signals to MDM4.

Besides MDM4, the MDM2 protein is also a target of stress signaling, and this is already known to include nucleolar stress. Not only the 5S RNP, including the ribosomal proteins L5 and L11, binds to MDM2 and prevents p53 ubiquitination (Donati et al., 2013; Onofrillo et al., 2017); other nucleolar proteins, such as nucleophosmin and p14^ARF^, move from nucleoli to the nucleoplasm upon nucleolar stress, where they bind and antagonize MDM2 (Kurki et al., 2004; Llanos et al., 2001; Sherr, 2006; Yang et al., 2016). Besides the 5S RNP, additional ribosomal proteins were found in association with MDM2, including L22 (Cao et al., 2017; Liu et al., 2016). Thus, L22 may activate p53 in two ways, by triggering *MDM4* exon skipping and by directly binding MDM2. Impaired activation of p53 in the absence of L22-binding RNA motifs within *MDM4* intron 6, as observed in our study, emphasizes the importance of the interaction between L22 and *MDM4* pre-mRNA.

Strikingly, in a genome-wide search for associations between exon expression levels and gene mutations in cancers, the link between *MDM4* exon 6 inclusion and deletions in the L22-coding gene was one of the top hits (Ghandi et al., 2019). Our results provide a mechanistic explanation for this; when L22 levels are lowered by deletion of at least one *L22* allele, this presumably prevents L22-induced exon skipping of *MDM4* and thus increases the inclusion of exon 6. Even more remarkably, gene losses of *L22* are particularly found in endometrial carcinomas of the microsatellite instability (MSI) subtype (Novetsky et al., 2013). This subtype often retains wildtype *TP53* (Schultheis et al., 2016). In this situation, it is conceivable that elevating full-length MDM4 through *L22* deletions will diminish p53 activity and thus allow enhanced tumor cell proliferation. L22 may thus be considered a haploinsufficient tumor suppressor, based on the tumor-promoting traits induced by deleting one of its alleles. L22 is not only unique in this regard, but it is also the only dispensable ribosomal protein, due to the existence of the paralogue L22L1. Indeed, the targeted deletion of *L22* does not preclude the survival of mice due to back-up by L22L1 (O’Leary et al., 2013).

Beyond *MDM4*, our study highlights two additional pre-mRNA targets of L22, the *L22L1* and *UBAP2L* pre-mRNAs. A broader spectrum of splicing regulation is consistent with a previously reported role of both L22 and L22L1 in zebrafish embryos; there, the ribosomal proteins affect the splicing of *smad2* and *smad1* in opposite directions (Zhang et al., 2013; Zhang et al., 2017).

To maintain the total levels of L22/L22L1, the negative control of *L22L1* expression by L22 appears to be required. The ability of L22 to bind the *L22L1* pre-mRNA and suppress its expression was reported previously (O’Leary et al., 2013), but the precise impact of L22 on the *L22L1* splicing pattern is first described here.

The other newly identified L22 splicing target, UBAP2L (aka NICE4), is essential for RNA polymerase II degradation when the enzyme hits a DNA lesion (Herlihy et al., 2022). Moreover, it is a core component of stress granules that modulate mRNA processing and storage under various stress conditions (Asano-Inami et al., 2023; Cirillo et al., 2020). Interestingly, UBAP2L is an RNA-binding protein that regulates the expression of genes that determine the global level of translation, such as *EIF4G1* (Luo et al., 2020). It is thus tempting to speculate that L22-mediated suppression of UBAP2L might serve to adapt global translational activity to the availability of ribosomes.

L22 was first discovered as a target of viral RNAs, especially the EBERs, i.e. highly abundant transcripts of RNA polymerase III, encoded by Epstein-Barr virus (Toczyski et al., 1994). It is still unknown which evolutionary advantage led viruses to express L22-interacting RNAs, but one possibility is that this might allow them to modulate the p53-regulatory system. By competing with the binding of L22 to *MDM4* pre-mRNA, the virus might increase the levels of full-length MDM4 and thus attenuate the p53 response.

According to our results, L22 connects three layers of gene expression: biogenesis of the translational machinery, mRNA maturation via pre-mRNA splicing, and RNA synthesis by transcription. Dysfunctional production of the ribosomal subunits triggers an extra-ribosomal splicing-regulatory function of L22. Subsequently, L22-driven alternative splicing of *MDM4* determines the activity of p53, the most prominent transcriptional regulator in the context of tumor suppression.

## ACKNOWLEDGEMENTS

We thank Manuel Kaulich for kindly sharing the RPE Cas9 cell line with us, and Dirk Görlich for providing the RNase inhibitor RNasin. We thank Antje Dickmanns, Sabrina Gerber and Xin Wang for helpful advice on methodology, and May-Britt Decker and Helene Dietrich for technical support. This work was supported by the Deutsche Forschungsgemeinschaft (DFG) via SFB1565, project number 469281184 (P12 to K.E.B., P06 to M.T.B. and P08 to M.D.). J.J. was supported by a doctoral stipend provided by the Studienstiftung des deutschen Volkes, and by the Göttingen Graduate School for Neurosciences, Biophysics, and Molecular Biosciences (GGNB).

## AUTHOR CONTRIBUTIONS

Conceptualization, J.J. and M.D.; methodology, J.J.; validation, J.J., K.E.B. and S.B.-F.; investigation, J.J.; writing – original draft, M.D. and J.J.; writing – review and editing, J.J., K.E.B., M.T.B. and M.D.; supervision, M.D. and M.T.B.

## MATERIAL AND METHODS

### Antibodies

| Antibody | Source | Identifier |
| --- | --- | --- |
| Alexa Fluor 488 goat anti-rabbit | Thermo Fisher Scientific | Cat# A-11034; RRID: AB_2576217 |
| Donkey anti-mouse IgG, HRP conjugated | Jackson ImmunoResearch Labs | Cat# 715-036-150; RRID: AB_2340773 |
| Donkey anti-rabbit IgG, HRP conjugated | Jackson ImmunoResearch Labs | Cat# 711-036-152; RRID: AB_2340590 |
| DYKDDDDK Tag (IF) | Cell Signaling | Cat# 14793; RRID: AB_2572291 |
| DYKDDDDK Tag (WB) | GenScript | Cat# A00187; RRID: AB_1720813 |
| FLAG M2-Peroxidase (HRP) (WB) | Sigma-Aldrich | Cat# A8592; RRID: AB_439702 |
| GAPDH (WB) | Abcam | Cat# ab8245; RRID: AB_2107448 |
| MDM2 IF2 (WB) | Millipore | Cat# OP46-100UG; RRID: AB_564803 |
| MDM4 (WB) | Millipore | Cat# 04-1555; RRID: AB_10562652 |
| p21 (WB) | Cell Signaling Technology | Cat# 2947; RRID: AB_823586 |
| p53 DO-1 (WB) | Santa Cruz Biotechnology | Cat# sc-126; RRID: AB_628082 |
| L11 (WB) | Cell Signaling Technology | Cat# 18163; RRID: AB_2798794 |
| L22 (IF) | Proteintech | Cat# 25002-1-AP; RRID: AB_2879841 |
| L22 (WB) | Santa Cruz Biotechnology | Cat# sc-136413; RRID: AB_10658965 |
| L5 (WB) | Cell Signaling Technology | Cat# 51345; RRID: AB_2799391 |
| S13 (WB) | Santa Cruz Biotechnology | Cat# sc-398690 |
| UBAP2L (WB) | Cell Signaling Technology | Cat# 40199 |
| ZMAT3 (WB) | Proteintech | Cat# 10504-1-AP; RRID: AB_2217579 |
| β-actin (WB) | Abcam | Cat# ab6276; RRID: AB_2223210 |

### Chemicals, peptides, and recombinant proteins

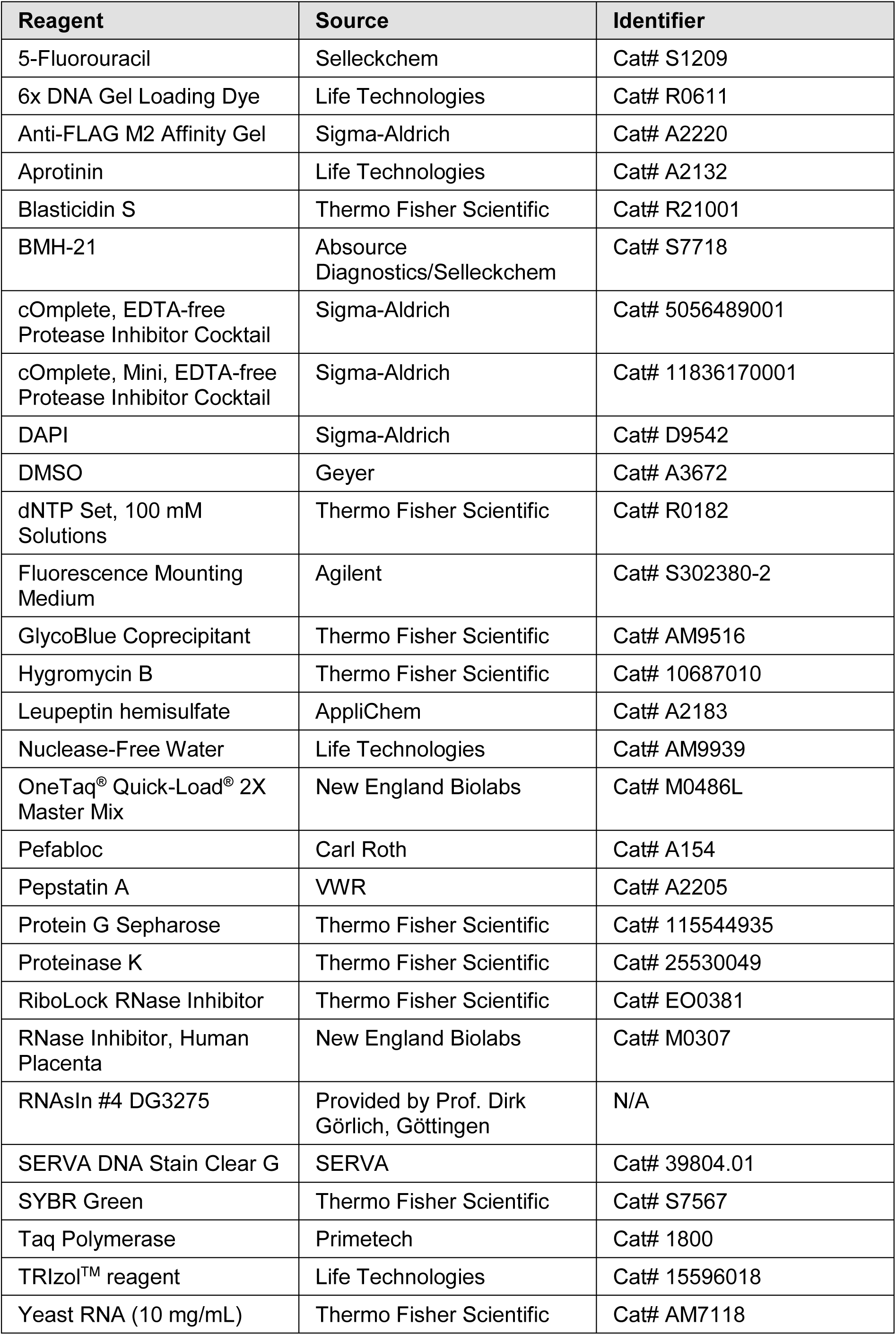

### Critical commercial assays

| Assay | Source | Identifier |
| --- | --- | --- |
| Ambion RNase III | Thermo Fisher Scientific | Cat# AM2290 |
| DNase I, RNase-free | Thermo Fisher Scientific | Cat# EN0521 |
| Immobilon Western HRP Substrate | Geyer/Millipore | Cat# WBKLS0500 |
| Lipofectamine 3000 transfection reagent | Thermo Fisher Scientific | Cat# L3000015 |
| MinElute PCR Purification Kit | Qiagen | Cat# 28006X4 |
| M-MuLV Reverse Transcriptase | New England Biolabs | Cat# M0253 |
| Pierce BCA Protein Assay Kit | Thermo Fisher Scientific | Cat# 23227 |
| QuikChange XL Site-Directed Mutagenesis Kit | Agilent | Cat# 200516 |
| SuperSignal West Femto Maximum Sensitivity Substrate | Thermo Fisher Scientific | Cat# 34095 |
| X-tremeGENE HP DNA Transfection Reagent | Sigma-Aldrich/Roche | Cat# 6366546001 |

### Experimental models: Cell lines

| Cell line | Source | Identifier |
| --- | --- | --- |
| Human: H1299 | ATCC | Cat# CRL-5803; RRID: CVCL_0060 |
| Human: HEK293 Flp-In T-REx | Thermo Fisher Scientific | Cat# R78007; RRID: CVCL_U427 |
| Human: HEK293 Flp-In L22-FLAG | This study | N/A |
| Human: HEK293T | ATCC | Cat# CRL-3216; RRID: CVCL_0063 |
| Human: RPE | ATCC | Cat# CRL-4000; RRID: CVCL_4388 |
| Human: RPE Cas9 | Provided by Prof. Manuel Kaulich, Frankfurt | N/A |
| Human: RPE *MDM4* $\Delta$ intron 6 | This study | N/A |

### Oligonucleotides

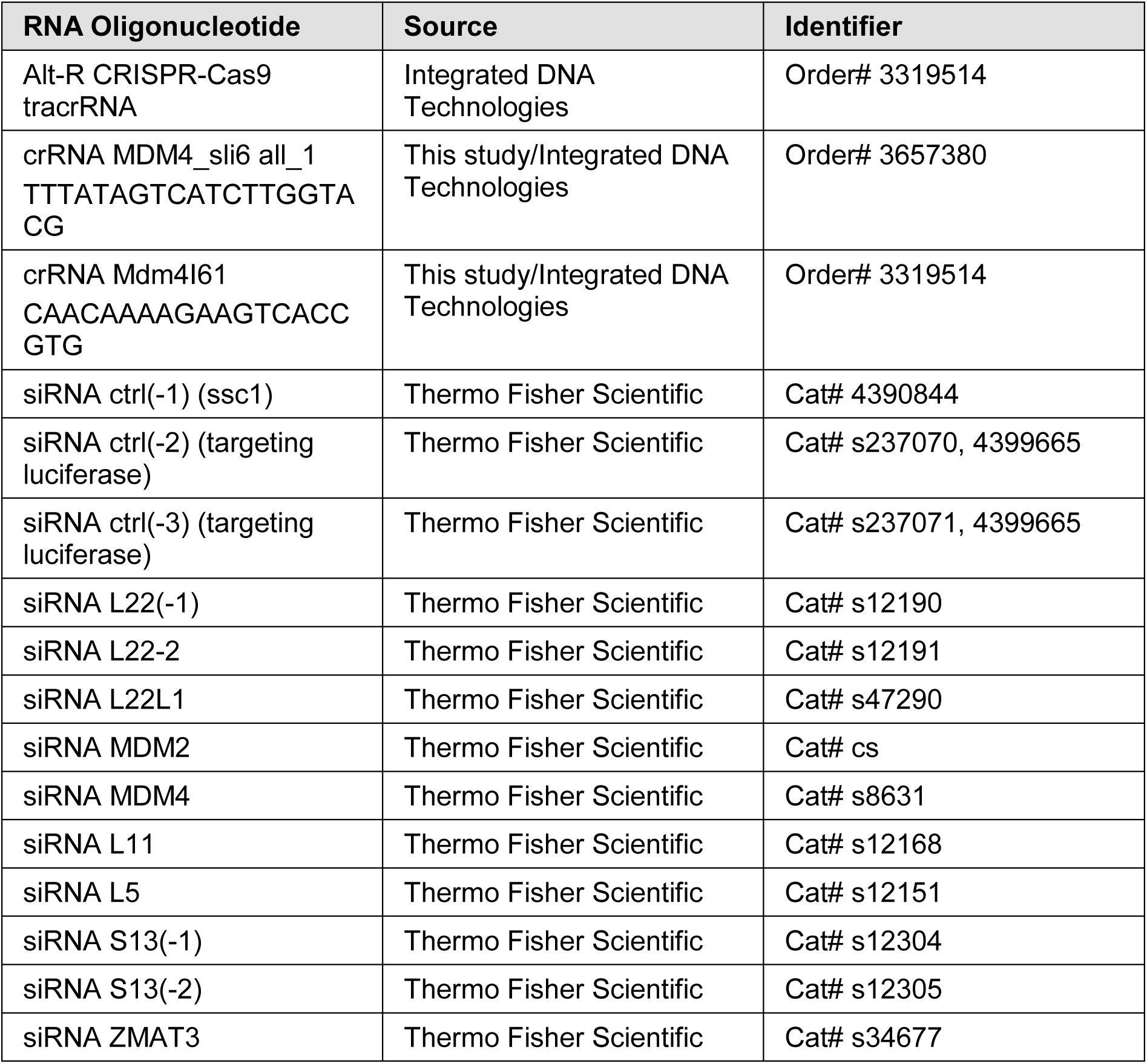

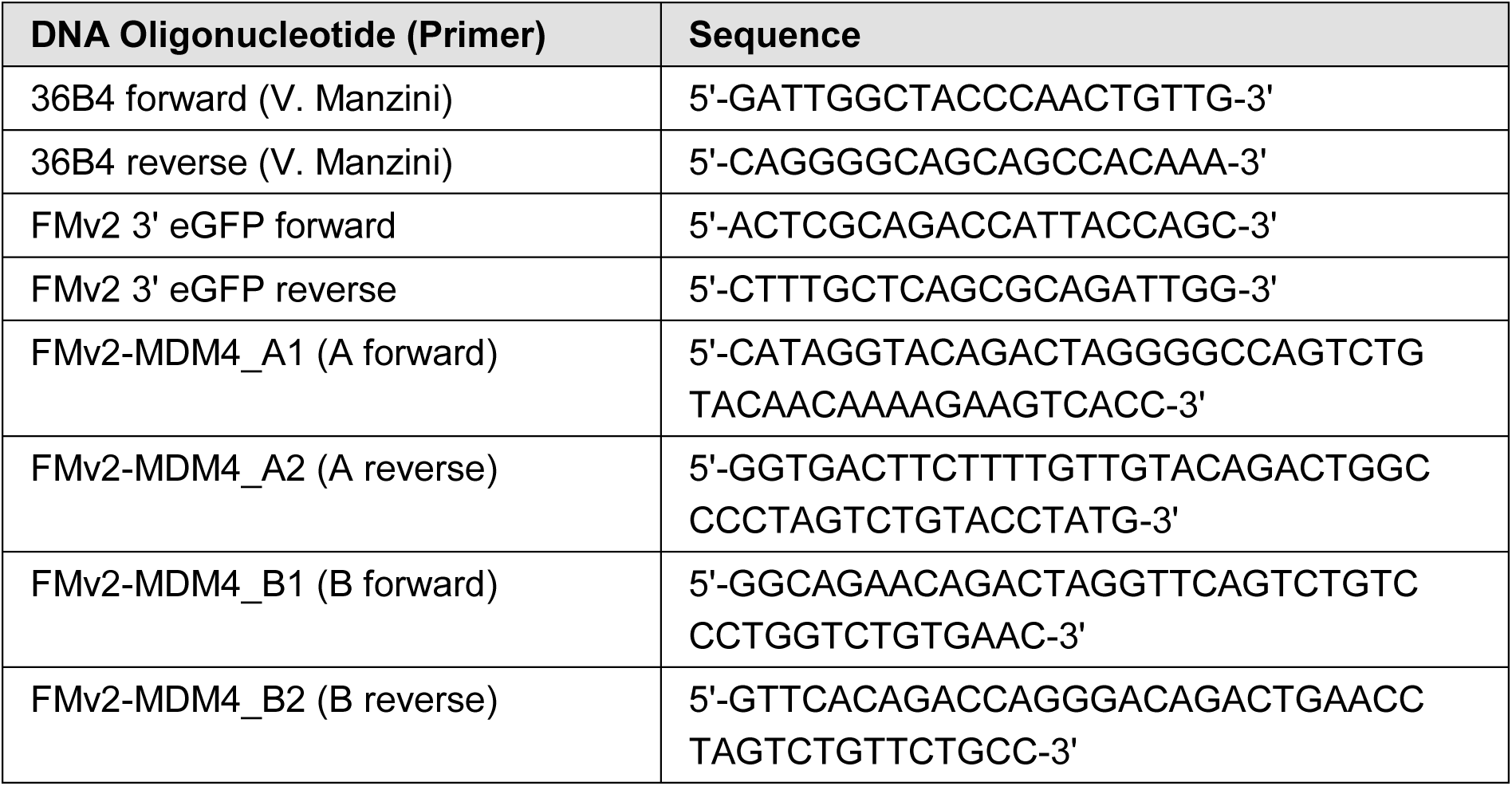
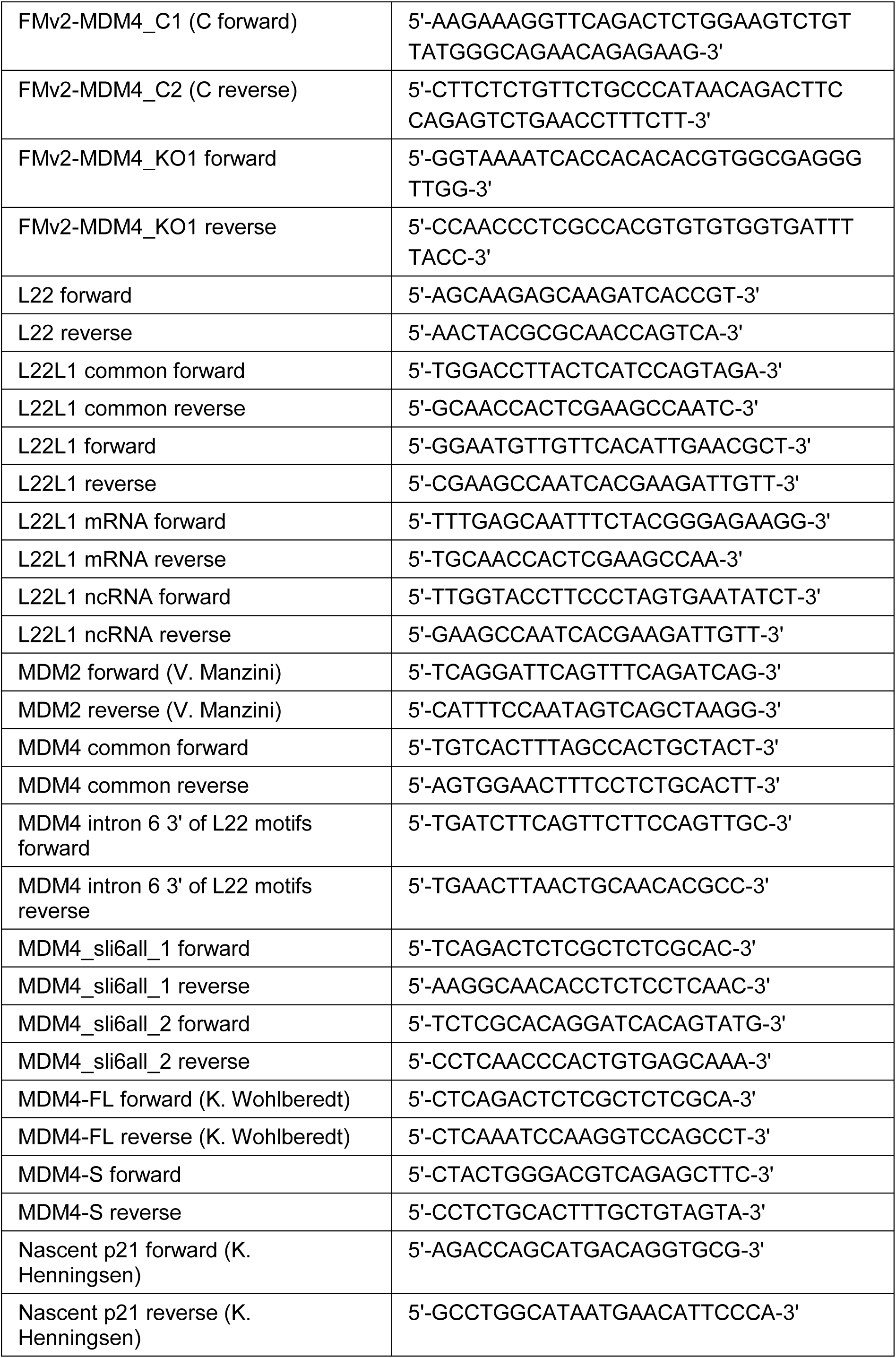
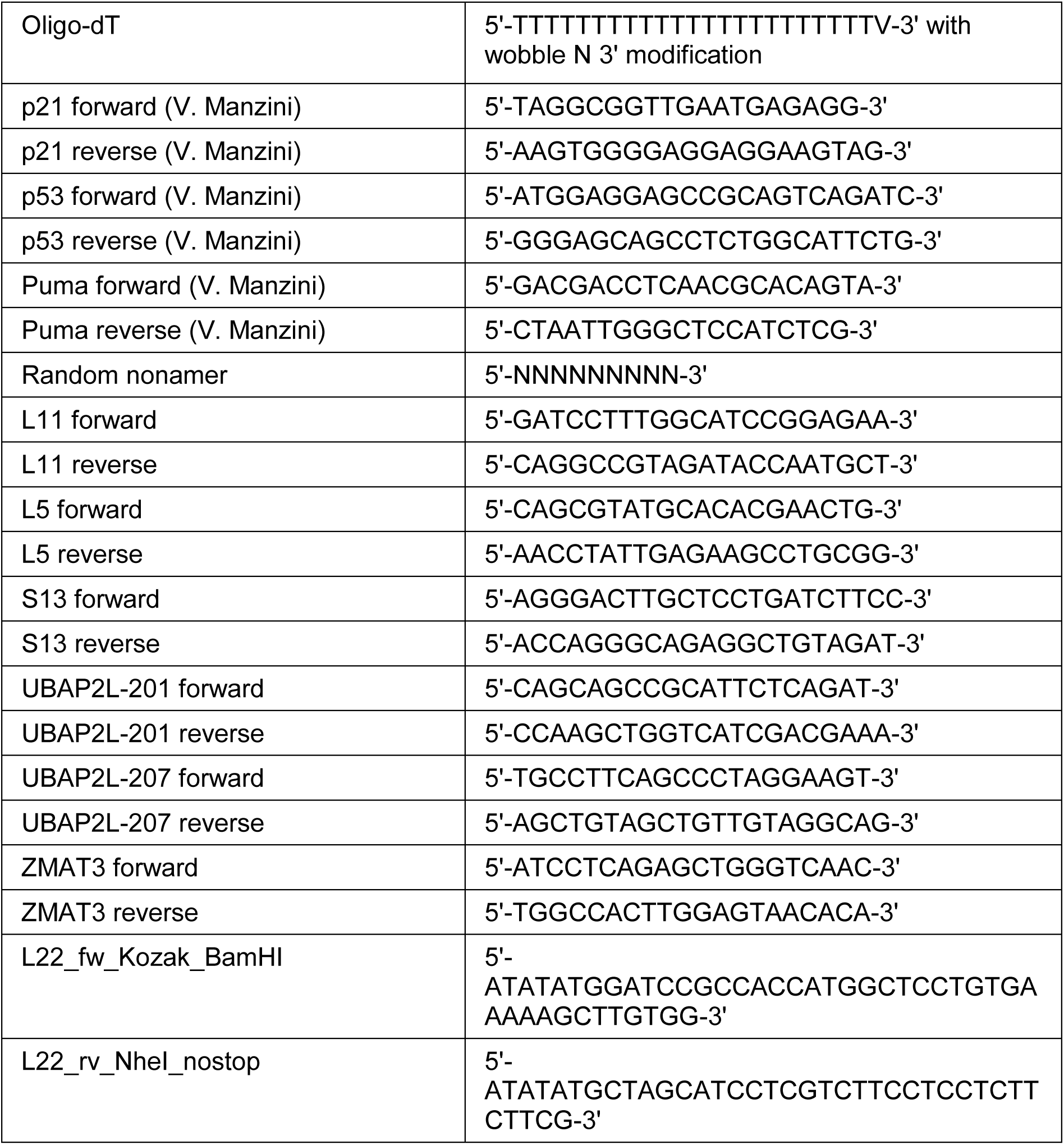

### Recombinant DNA

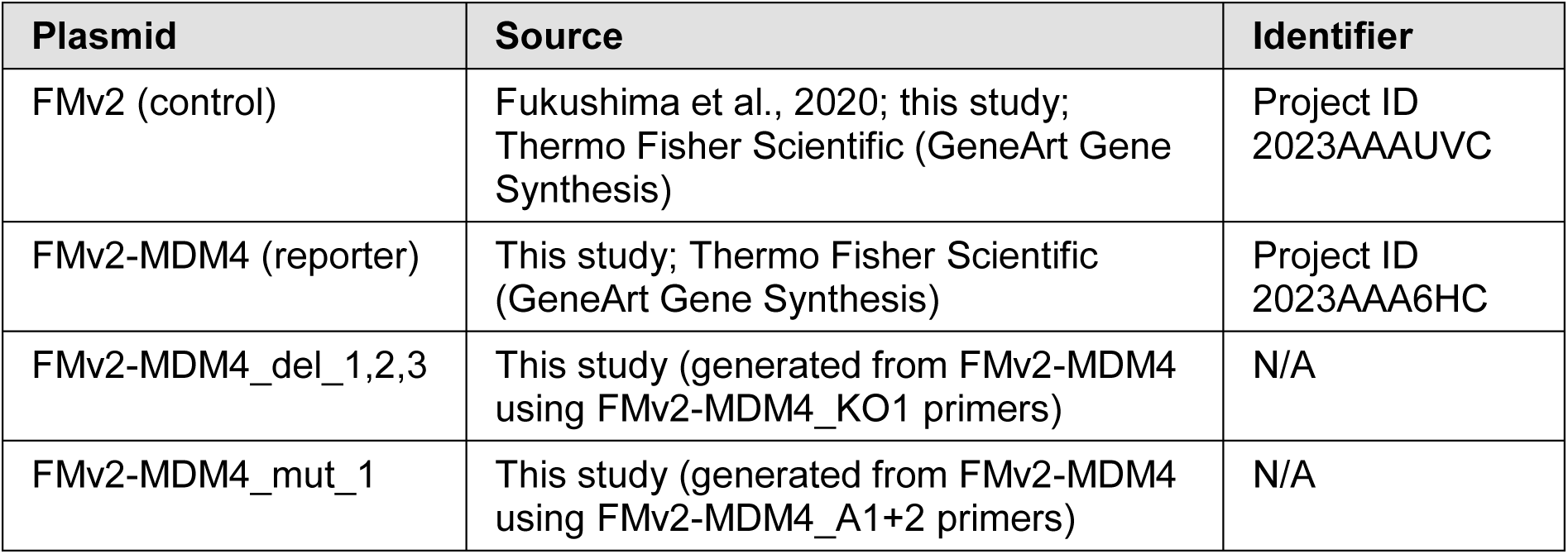
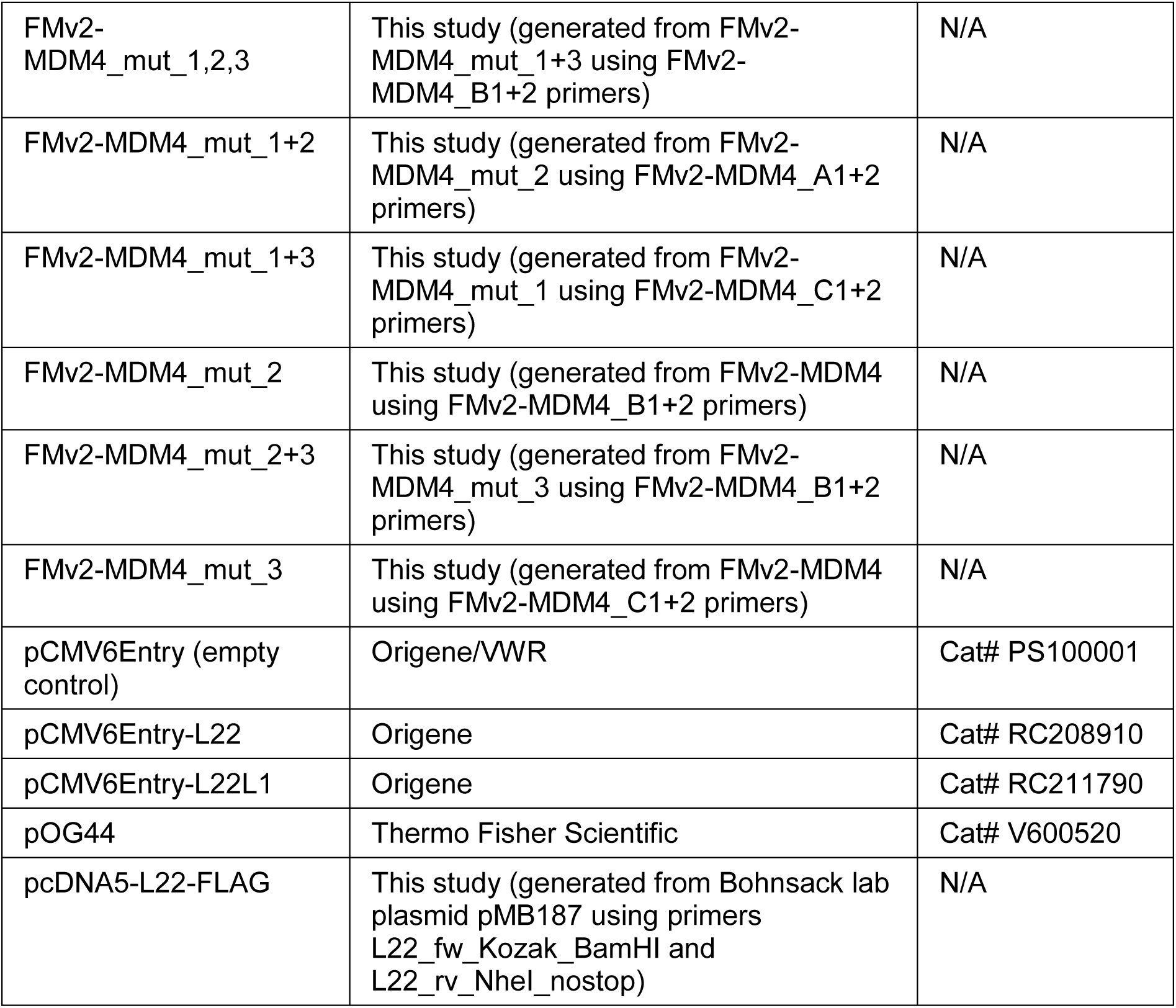

| Plasmid | Source | Identifier |
| --- | --- | --- |
| FMv2 (control) | Fukushima et al., 2020; this study; Thermo Fisher Scientific (GeneArt Gene Synthesis) | Project ID 2023AAAUVC |
| FMv2-MDM4 (reporter) | This study; Thermo Fisher Scientific (GeneArt Gene Synthesis) | Project ID 2023AAA6HC |
| FMv2-MDM4_del_1,2,3 | This study (generated from FMv2-MDM4 using FMv2-MDM4_KO1 primers) | N/A |
| FMv2-MDM4_mut_1 | This study (generated from FMv2-MDM4 using FMv2-MDM4_A1+2 primers) | N/A |
| FMv2-MDM4_mut_1,2,3 | This study (generated from FMv2-MDM4_mut_1+3 using FMv2-MDM4_B1+2 primers) | N/A |
| FMv2-MDM4_mut_1+2 | This study (generated from FMv2-MDM4_mut_2 using FMv2-MDM4_A1+2 primers) | N/A |
| FMv2-MDM4_mut_1+3 | This study (generated from FMv2-MDM4_mut_1 using FMv2-MDM4_C1+2 primers) | N/A |
| FMv2-MDM4_mut_2 | This study (generated from FMv2-MDM4 using FMv2-MDM4_B1+2 primers) | N/A |
| FMv2-MDM4_mut_2+3 | This study (generated from FMv2-MDM4_mut_3 using FMv2-MDM4_B1+2 primers) | N/A |
| FMv2-MDM4_mut_3 | This study (generated from FMv2-MDM4 using FMv2-MDM4_C1+2 primers) | N/A |
| pCMV6Entry (empty control) | Origene/VWR | Cat# PS100001 |
| pCMV6Entry-L22 | Origene | Cat# RC208910 |
| pCMV6Entry-L22L1 | Origene | Cat# RC211790 |
| pOG44 | Thermo Fisher Scientific | Cat# V600520 |
| pcDNA5-L22-FLAG | This study (generated from Bohnsack lab plasmid pMB187 using primers L22_fw_Kozak_BamHI and L22_rv_NheI_nostop) | N/A |

### Software and algorithms

| Software and algorithms | Source | Identifier |
| --- | --- | --- |
| BioRender | BioRender | https://app.biorender.com |
| Fiji v. 2.0.0 | NIH | N/A |
| Image Lab v. 5.2.1 | Bio Rad | N/A |
| Prism v. 9.3.1 | GraphPad | N/A |
| RNAfold web server | Institute for Theoretical Chemistry, University of Vienna | http://rna.tbi.univie.ac.at/cgi-bin/RNAWebSuite/RNAfold.cgi |

### Cell culture

The human non-small cell lung cancer cell line H1299 and the human embryonic kidney cell line containing the SV40 T-antigen HEK293T were maintained in Dulbeccós Modified Eaglés Medium (DMEM, 31600091, Thermo Fisher Scientific) supplemented with 10% Fetal Bovine Serum (FBS) (ACSM0190, Anprotec), 2 mM glutamine (25030123, Thermo Fisher Scientific), 50 U/mL penicillin, 50 µg/mL streptomycin (15140122, GIBCO), 10 µg/mL ciprofloxacin (PZN03277618, Fresenius Kabi), and 2 µg/mL tetracycline (87128, Sigma-Aldrich). To generate a stably transfected cell line for the inducible expression of L22 with a C-terminal FLAG tag, HEK293 Flp-In T-Rex cells were reverse transfected with an appropriate pcDNA5-based plasmid and pOG44, which expresses the Flp recombinase, using X-tremeGENE HP DNA transfection reagent (6366546001, Sigma-Aldrich/Roche) according to the manufacturer’s instructions. Cells with appropriate genomic integrations in the Flp-In locus were selected using 100 µg/mL hygromycin B (10687010, Thermo Fisher Scientific) and 10 µg/mL blasticidin S (R21001, Thermo Fisher Scientific) for approximately two weeks. Colonies were resuspended, pooled and cells were maintained in DMEM GlutaMAX (61965059, Gibco) supplemented with 10% FBS, penicillin, streptomycin, and 100 µg/mL hygromycin B. Additionally, 10 µg/mL blasticidin S was added to the culture medium once a week. The selection markers hygromycin B and blasticidin S were removed from culturing medium 24 h prior to experiments. Expression of L22 was induced by adding 1 µg/mL tetracycline for 24 h to 48 h. The non-tumorigenic human retinal pigment epithelial cell line RPE and its derivatives RPE p53^-/-^ (Klusmann et al., 2016), RPE Cas9 and RPE *MDM4* Δ intron 6 were cultured in DMEM GlutaMAX supplemented with 10% FBS, penicillin, streptomycin, ciprofloxacin, and 50 µg/mL hygromycin B for hTERT activation. Hygromycin B was removed from culturing medium 24 h prior to experiments. All cell lines were cultured at 37 °C with 5% CO_2_ and routinely tested for mycoplasma contamination.

Cells were treated with BMH-21 (S7718, Absource Diagnostics/Selleckchem), 5-fluorouracil (S1209, Selleckchem), and DMSO (A3672, Geyer).

Transfections of siRNAs were performed using Lipofectamine 3000 (L3000015, Thermo Fisher Scientific). Cells were reverse transfected with final siRNA concentration of 12.5 nM siRNA. Medium was refreshed after 24 h, and cells were harvested after 48 h or 72 h.

Cells were forward transfected with plasmids in amounts of 2.5 µg per well of 6-well plates Medium was refreshed after 6 h (except for HEK293-derived cell lines, to avoid cell detachment), and cells were harvested after 42 h or 48 h.

### CRISPR/Cas9-mediated generation of knockout cells

To delete all three L22 consensus sequences in *MDM4* intron 6, crRNAs binding upstream (Mdm4I61) and downstream (MDM4_sli6all_1) of these sequences were designed using the web tool CRISPOR.org (Concordet and Haeussler, 2018). For crRNA:tracrRNA (Integrated DNA Technologies) 50 µM duplex formation, equal volumes of 100 µM crRNA and 100 µM tracrRNA were mixed, incubated at 95 °C for 5 min and cooled to room temperature.

crRNA:tracrRNA duplexes were transfected into RPE Cas9 cells by reverse transfection using Lipofectamine 3000 transfection reagent (L3000015, Thermo Fisher Scientific) with a final crRNA:tracrRNA concentration of 25 nM (12.5 nM upstream crRNA:tracrRNA, 12.5 nM downstream cr:tracrRNA). Medium was refreshed after 24 h.

At 48 h post transfection, cells were trypsinized and 50, five or two cells per well were seeded on 96-well plates. After around three weeks, cells from different wells were passaged further to larger plates. RPE *MDM4* Δ intron 6 cells, a cell line generated from the 96-well plate with 50 cells per well, were chosen for subsequent experiments since they showed a complete knockout in genotyping and sequencing.

### DNA isolation, genotyping, and sequencing

Genotyping and sequencing were performed to identify RPE cell lines generated by CRISPR/Cas9 with deletion of all three L22 consensus sequences in *MDM4* intron 6. For DNA isolation, a 2 mL aliquot of cell suspension was centrifuged briefly (3000 rpm, 5 min, 4 °C) and pellets were washed once with PBS. Each pellet was resuspended in 50 µL 0.1% SDS shearing buffer (1% Triton X-100, 0.15 M NaCl, 1 mM EDTA, 0.5 mM EGTA, 20 mM HEPES, 0.1% SDS), followed by the addition of 50 µL 10 mM Tris, 100 µL 2x DNA isolation buffer pH 8.0 (100 mM Tris-HCl pH 8.8, 20 mM EDTA, 2% SDS) and 2 µL 20 mg/mL proteinase K (25530049, Thermo Fisher Scientific). Samples were incubated overnight at 65 °C and 800 rpm, 200 µL 10 mM Tris was added, and DNA was isolated using the MinElute PCR Purification Kit (28006X4, Qiagen).

For genotyping, MDM4_sli6all_1 primers (Metabion) binding upstream and downstream of the crRNA binding sites were used. One genotyping PCR sample contained 10 µL OneTaq^®^ Quick-Load^®^ 2X Master Mix (M0486L, New England Biolabs), 1 µL 10 µM forward primer, 1 µL 10 µM reverse primer, 3 µL ddH_2_O, and 5 µL DNA. The PCR program was a preheating step (2 min, 95 °C), followed by 36 cycles of 95 °C for 30 sec, 56 °C for 1 min and 72 °C for 1.5 min, with a final step at 72 °C for 5 min. Genotyping PCR products were separated by agarose gel electrophoresis using SERVA DNA Stain Clear G (39804.01, SERVA) for DNA detection under a UV illuminator.

DNA samples showing the PCR product sizes expected for successful knockout in genotyping were sequenced for further confirmation. For this, genotyping PCR products were purified using MinElute PCR Purification Kit (28006X4, Qiagen), DNA concentration was measured by spectrophotometry, and samples were sequenced with MDM4_sli6all_2 forward and reverse primers (Eurofins Genomics).

### RNA isolation, reverse transcription, and real-time quantitative PCR

Cells were washed with PBS and harvested in TRIzol^TM^ reagent containing phenol (15596018, Life Technologies). For total RNA extraction, 200 µL chloroform per 1 mL TRIzol^TM^ was added. Samples were centrifuged for phase separation, and the upper aqueous phase was used for precipitating the RNA by adding 500 µL isopropanol – and optionally 2 µL GlycoBlue Coprecipitant (AM9516, Thermo Fisher Scientific) – and centrifuging. The RNA pellet was washed once with 75% ethanol, dried, and resuspended in nuclease-free H_2_O (AM9939, Life Technologies). RNA concentration was measured by spectrophotometry.

For reverse transcription, 1 µg total RNA was incubated with 6.25 µM oligo-dT (Metabion) and 1.875 µM random nonamer primers (Metabion) as well as 0.625 mM dNTPs (R0182, Thermo Fisher Scientific) at 70 °C for 5 min for primer annealing. Next, 10 U of RNase Inhibitor, Human Placenta (M0307, New England Biolabs) and 25 U of M-MuLV Reverse Transcriptase with corresponding buffer (M0253, New England Biolabs) was added and the reaction was incubated at 42 °C for 1 h, followed by 5 min incubation at 95 °C for inactivating the reverse transcriptase. A control reaction without reverse transcriptase (no reverse transcriptase control, NRT) was included. Finally, the synthesized cDNA was diluted by adding 130 µL ddH_2_O per 20 µL cDNA reaction mixture and used as a template for real-time quantitative PCR (RT-qPCR).

For quantification, RT-qPCR was performed in technical duplicates. One well of a 96-well plate contained one reaction consisting of 1 µL 10 µM forward primer, 1 µL 10 µM reverse primer, 3 µL cDNA, 6 µL ddH_2_O, and 14 µL of a homemade RT-qPCR reaction mixture containing 518 mM trehalose (stock in 10 mM Tris pH 8.0), 130 mM Tris-HCl pH 8.8, 35 mM ammonium sulphate, 0.017% Tween-20, 0.43% Triton X-100, 5.18 mM MgCl_2_, 0.43x concentrated SYBR Green (S7567, Life Technologies), 0.35 mM dNTPs (R0182, Thermo Fisher Scientific), and 35 U/mL Taq polymerase (1800, Primetech). The program used for the thermocycler was a preheating step (2 min, 95 °C), followed by 45 cycles of 95 °C for 15 sec and 60 °C for 1 min. Subsequently, a melting curve was generated, followed by a final incubation at 95 °C for 30 sec.

Gene expression levels were normalized to the mRNA levels of the housekeeping gene *36B4*, and the analysis was conducted using the ΔΔCt method.

For semi-quantitative detection of *MDM4* and *L22L1* splice variants using common primers leading to products of different sizes for the respective splice variants, RT-qPCR was performed as usual. Next, 6x DNA Gel Loading Dye (R0611, Life Technologies) was added to RT-qPCR products and they were separated and visualized by agarose gel electrophoresis using SERVA DNA Stain Clear G (39804.01, SERVA) for DNA detection under a UV illuminator. Band quantification was performed using ImageJ. Each band was defined as a region of interest (ROI) and a background ROI was defined as well. Band intensities were determined and resulting “mean” values of sample minus background were used to determine splice variant ratios.

### Immunoblot analysis

Cells were washed with PBS and harvested in lysis buffer, i.e. 14 mM Tris-HCl pH 7.5, 100 mM NaCl, 9 mM EDTA, 1% Triton X-100, 1% deoxycholate, 0.1% SDS, 2.7 M urea, supplemented with protease inhibitors (aprotinin, leupeptin hemisulfate, aminoethyl-benzene-sulfonyl fluoride/Pefabloc, pepstatin A). The samples were briefly sonicated and pushed through a syringe to disrupt chromatin. Protein concentration was measured using a BCA protein assay kit (23227, Thermo Fisher Scientific). Equal protein amounts of each sample were boiled in Laemmli buffer at 95 °C for 5 min and separated by SDS-PAGE. Proteins were transferred to a nitrocellulose membrane. Membranes were blocked with 5% (w/v) non-fat milk in TBS containing 0.1% Tween-20 for 1 h and incubated with the primary antibodies at 4 °C overnight. Next, membranes were incubated with horse radish peroxidase (HRP)-conjugated secondary antibodies (donkey anti-mouse or anti-rabbit IgG, Jackson ImmunoResearch Labs), followed by protein detection using either Immobilon Western HRP Substrate (WBKLS0500, Geyer/Millipore) or SuperSignal West Femto Maximum Sensitivity Substrate (34095, Thermo Fisher Scientific).

### RNA immunoprecipitation (RIP)

For RIP to precipitate FLAG-tagged L22 and its bound RNAs, cells grown on six wells of 6-well plates per sample were washed once with PBS, and RNA and protein were crosslinked with 0.75% paraformaldehyde (PFA) in PBS for 10 min. Fixation was quenched by adding 0.125 M glycine for 5 min. Next, cells were washed once with PBS and harvested on ice in RIP high salt buffer (500 mM NaCl, 50 mM Tris, 5 mM EDTA, 0.5% NP-40), supplemented with protease inhibitors (11836170001, Sigma-Aldrich) and, in case of samples used not only for protein but also for bound RNA precipitation, with 0.1 U/µL RNase inhibitor (RNAsIn #4 DG3275, kindly provided by D. Görlich). All following steps were performed on ice. Samples were pushed through a syringe and sonicated to disrupt chromatin, followed by centrifugation for 10 min at 16100 rcf and 4 °C to remove cell debris, proceeding with the supernatant. The cell lysate was pre-cleared with Protein G Sepharose (115544935, Thermo Fisher Scientific) for 2 h at 4 °C, and 5% of each sample was preserved as input. This was followed by an overnight incubation at 4 °C with 30 µL (15 µL beads) per sample of Anti-FLAG M2 Affinity Gel (A2220, Sigma-Aldrich). Additionally, 20 µg/mL yeast RNA (AM7118, Thermo Fisher Scientific) was added for this precipitation step as a competitor RNA. Beads were washed three times with RIP high salt buffer and three times with RIP low salt buffer (5 mM NaCl, 50 mM Tris, 5 mM EDTA, 0.5% NP-40), supplemented with protease inhibitors and RNase inhibitor as described above. Samples only used for protein precipitation were washed once more with RIP high salt buffer. Beads were then resuspended in Laemmli buffer and boiled at 95 °C for 5 min prior to SDS-PAGE and immunoblot analysis using FLAG antibody covalently conjugated to horse radish peroxidase for detection of precipitated L22 (A8592, Sigma-Aldrich).

For samples also used for RNA precipitation, a digestion with RNase III (AM2290, Thermo Fisher Scientific) was performed to generate smaller fragments from precipitated RNAs. For this, beads were incubated with 0.05 U/µL RNase III and corresponding buffer at 37 °C for 30 min, followed by another washing step with RIP high salt buffer. The RNA-protein complexes of immunoprecipitation (IP) and input samples were then decrosslinked by adding 0.4 µg/µL proteinase K (25530049, Thermo Fisher Scientific) in RIP high salt buffer for 1 h at 60 °C. 500 µL TRIzol^TM^ reagent was added per sample to input and beads, and RNA was isolated as usual. Each RNA pellet was resuspended in 11 µL nuclease-free H_2_O, and DNase treatment was performed by incubating 8.8 µL RNA with 2 U DNase I, RNase-free with corresponding buffer (EN0521, Thermo Fisher Scientific) and 20 U RiboLock RNase Inhibitor (EO0381, Thermo Fisher Scientific) at 37 °C for 30 min. Next, 4.5 mM EDTA was added and samples were incubated at 65 °C for 10 min for DNase inactivation. 10 µL from 22 µL DNase-treated RNA was used as a template for reverse transcription as described above, except that RNA was incubated with 6.25 µM random nonamer primers (Metabion). A control reaction without reverse transcriptase (no reverse transcriptase control, NRT) was included for each sample. Finally, the synthesized cDNA was used as a template for RT-qPCR. The quantification of the RT-qPCR signal was performed relative to the input.

### Immunofluorescence

Cells were grown in 8-well chambers on PCA slides with removable frame (946140802, Sarstedt). They were fixed in 3.7% PFA in PBS for 30 min, rinsed with PBS, and permeabilized with 0.5% Triton X-100 in PBS for 30 min. After 2× 5 min washing with PBS, blocking solution (10% FBS 0.1% Tween-20 in PBS) was added to the cells for 10 min, followed by incubation with primary antibody produced in rabbit, diluted in blocking solution, at 4 °C overnight. Cells were washed 2× 5 min with PBS, and secondary antibody (Alexa Fluor 488 goat anti-rabbit; A-11034, Thermo Fisher Scientific) (1:500) and 4’,6-diamidino-2-phenylindole (DAPI; D9542, Sigma-Aldrich) (1:3000) diluted in blocking solution were added for 1 h at room temperature. Two more washing steps with PBS were performed, gaskets were removed and slides were mounted onto coverslips using Fluorescence Mounting Medium (S302380-2, Agilent). Images were acquired using a Zeiss Axio Scope.A1 fluorescence microscope with 40× and 100x magnifications and further analyzed with ImageJ. For quantification of nuclear L22, the DAPI channel was used to create binary images (segmentation) to define the nuclei as regions of interest (ROIs). These ROIs were then used to quantify the L22 signal within these regions. Mean fluorescence intensity values were visualized.

### *MDM4* splicing reporter plasmid generation

FMv2 (control plasmid) was generated by inserting the following sequences into the pcDNA3.1/Hygro(+) vector containing the CMV promoter using the restriction enzymes *Nhe*I and *Xho*I: the mCherry coding region, the 5′ end of the *eGFP* gene, the 5′ end of *DMD* intron 18, a Multiple Cloning Site (MCS), the 3′ end of *DMD* intron 19, and the 3′ end of the *eGFP* gene. FMv2-MDM4 (*MDM4* splicing reporter plasmid) was generated by inserting the following sequences into the MCS of FMv2 using the restriction enzymes *Hind*III and *Bam*HI: the 3′ end (34 bp) of *MDM4* intron 5, *MDM4* exon 6 (68 bp), and the 5′ end (398 bp) of *MDM4* intron 6 containing all three L22 consensus sequences. For both plasmids, the GeneArt Gene Synthesis service (Thermo Fisher Scientific) was used. Different FMv2-MDM4 versions containing mutations or deletions of single or combined L22 consensus sequences in *MDM4* intron 6 were generated by QuikChange site-directed mutagenesis (200516, Agilent).

### *MDM4* splicing reporter assay

The principle of this assay is as follows: upon overexpression of either FMv2 or FMv2-MDM4 or its mutated versions in cells, mCherry protein is constitutively expressed and serves as a transfection marker. eGFP, the sequence of which is split into two parts in all plasmids, is also constitutively expressed upon FMv2 overexpression, but only if the *MDM4* exon 6 is skipped in case of FMv2-MDM4 (or variant) overexpression, thus serving as a splicing marker.

For the assay, cells were grown on 24-well plates (3524, Corning Costar/Geyer), transfected with control (ctrl) or L22 (L22) siRNAs for 72 h, and with control plasmid (FMv2) (Fukushima et al., 2020), *MDM4* reporter plasmid (FMv2-MDM4) or different FMv2-MDM4 versions for 48 h. Using the ZEISS Celldiscoverer 7 live cell fluorescence microscope with 5x magnification, red fluorescence (mCherry, exposure time 100 ms, intensity 30%) and green fluorescence (eGFP, exposure time 20 ms, intensity 30%) were detected, and five to six images per well were taken.

Images were further analyzed with ImageJ. The mCherry channel was used to create binary images (segmentation) to define strongly transfected cells (i.e., cells with high mCherry intensity above a threshold) as regions of interest (ROIs). These ROIs were then used to quantify the eGFP signal within these regions. The average of all resulting “mean” values was calculated and the L22/ctrl ratio of these values was determined.

### Cell confluence measurements

Cells were plated on 24-well plates (3524, Corning Costar/Geyer) and continuously treated with the indicated compounds starting on the day after seeding, with a refreshment of treatment every other day. On each day of treatment, cell confluence was measured and analyzed using the Celigo Adherent Cell Cytometer.

### Statistical analysis

Statistical analysis was performed using GraphPad Prism software. Unpaired Student’s t test was used for statistical analysis of RT-qPCR, agarose gel band quantification, reporter experiments, and confluence measurements. Mann-Whitney U test was used for statistical analysis of quantification of single nuclear intensities of L22 immunofluorescence signal. Ns, not significant; *, P≤0.05; **, P≤0.01; ***, P≤0.001; ****, P≤0.0001.

## LEGENDS TO FIGURES

**Supplementary Figure 1:**
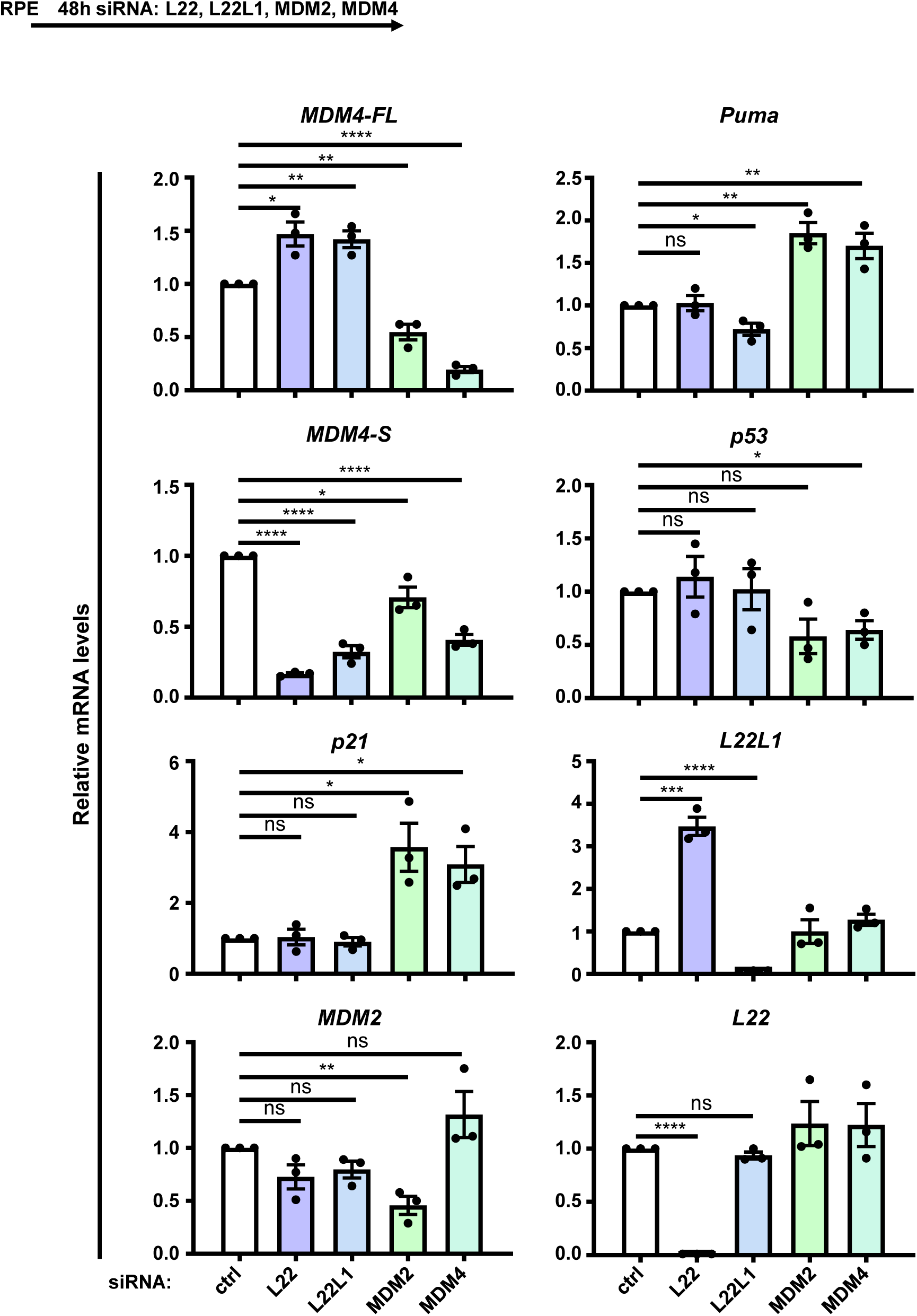
Gene expression upon depletion of L22 and L22L1, as well as MDM4 and MDM2. RPE cells were transfected with siRNAs to deplete L22, L22L1, MDM2, or MDM4, or with control (ctrl) siRNA, for 48 h. RT-qPCR analyses to detect the indicated target mRNAs were performed, with normalization to the *36B4* (housekeeping gene) mRNA level and then to the control sample. The average of three biological replicates is depicted. Error bars, standard error of the mean. The statistical significance was assessed by an unpaired t test: ns, not significant; *, P≤0.05; **, P≤0.01; ***, P≤0.001; ****, P≤0.0001.

**Supplementary Figure 2:**
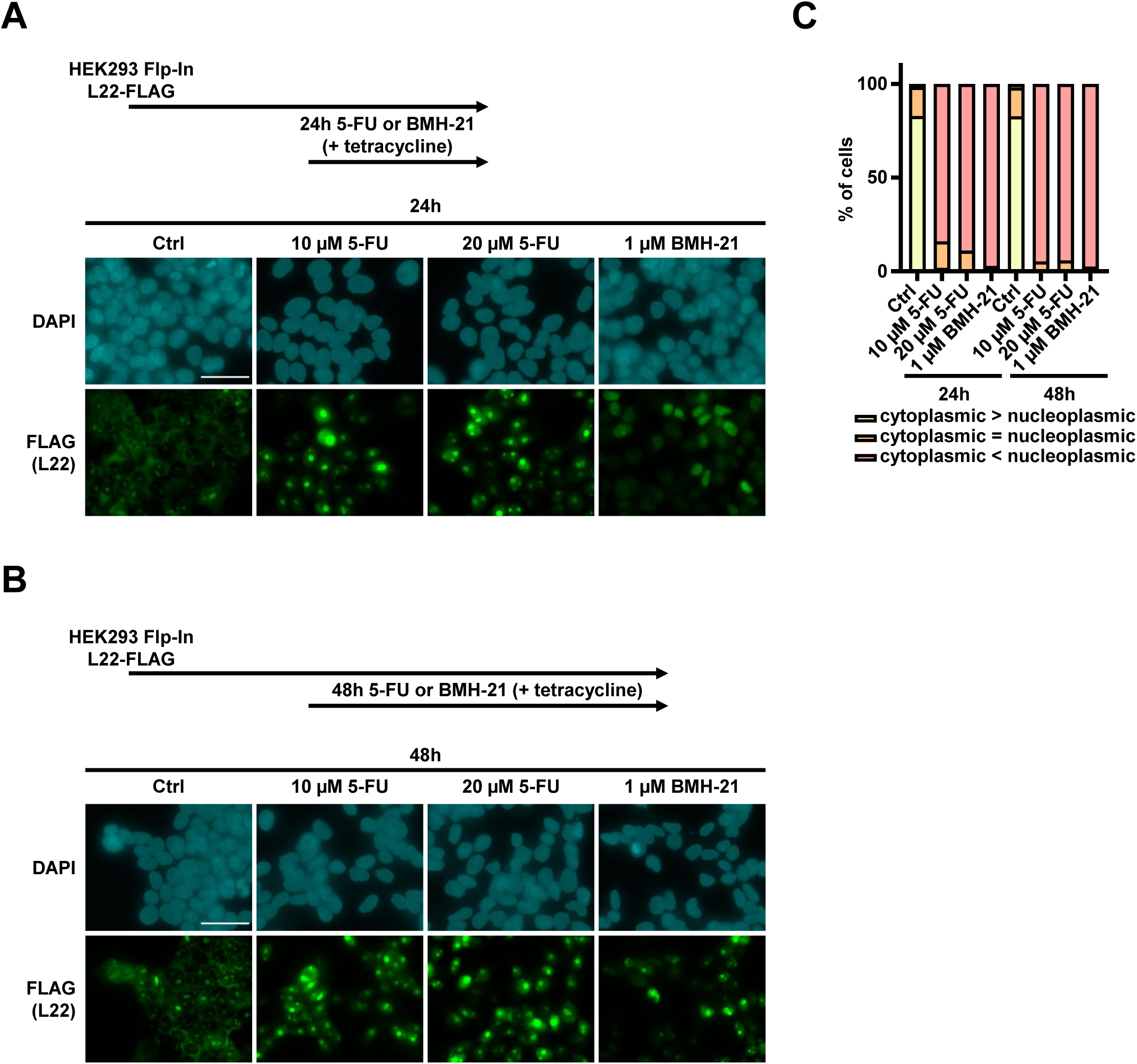
L22-FLAG accumulation in the nucleoplasm upon treatment with BMH-21 or 5-FU. **A.-C.** Stably transfected HEK293 Flp-In cells for the inducible expression of L22-FLAG were treated with 10 or 20 µM 5-FU or 1 µM BMH-21 or control-treated (ctrl), for 24 h or 48 h, to induce nucleolar stress. Tetracycline (1 µg/mL) was added for all conditions, including the control, during the entire treatment duration (24 h or 48 h), to induce the expression of FLAG-tagged L22. Immunofluorescence staining of L22 using a FLAG antibody was performed. **A.-B.** Representative images (100x objective, bar: 40 µm) of the nuclear (DAPI, blue) and L22 (green) staining upon 24 h (**A**) and 48 h treatment (**B**). **C.** Quantification of cytoplasmic versus nucleoplasmic intensity of the L22 signal. At least 100 cells were analyzed in each case.

**Supplementary Figure 3:**
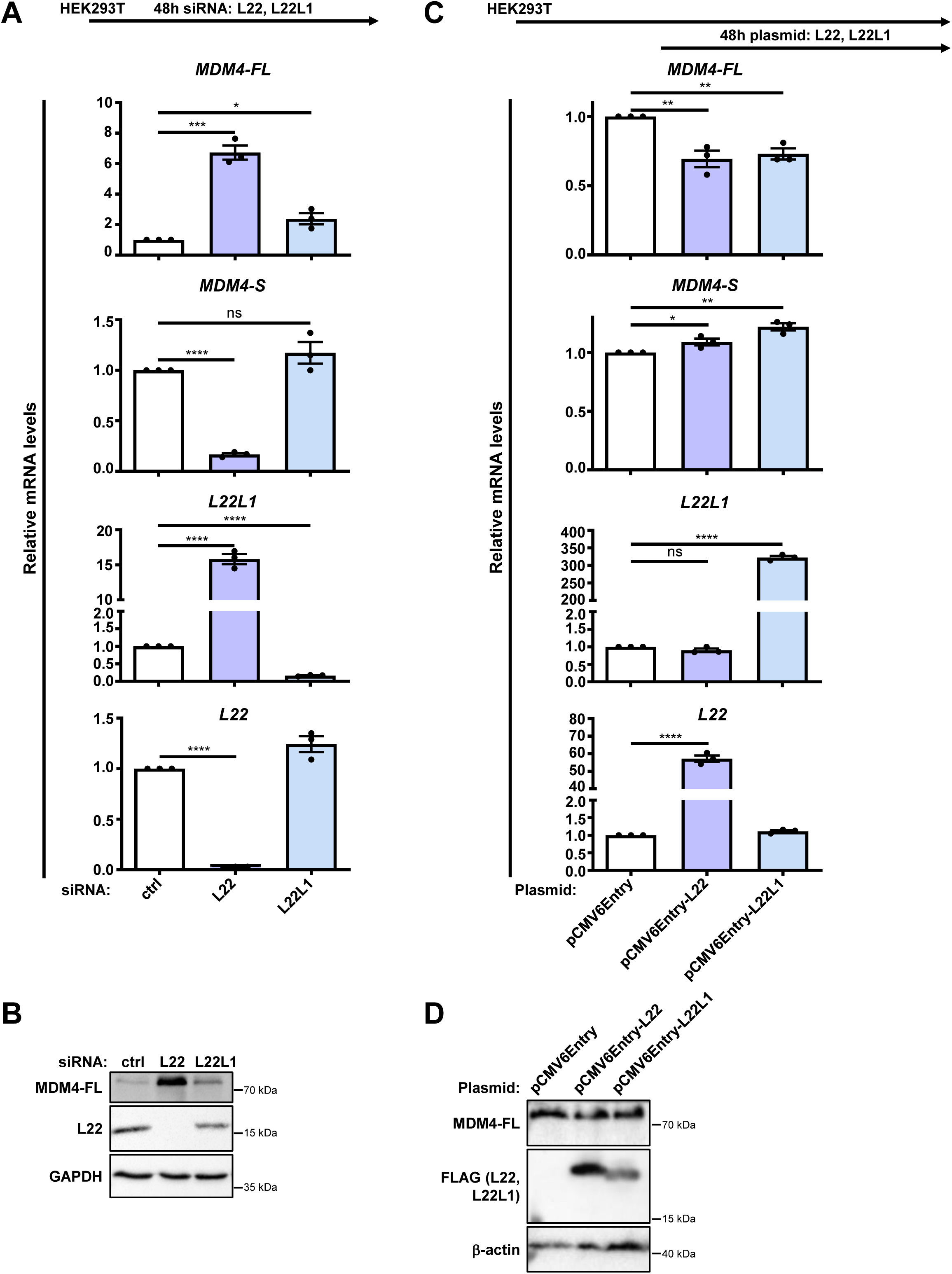
*MDM4* exon 6 inclusion upon depletion or overexpression of L22 or L22L1. **A.-B.** HEK293T cells were transfected with siRNA to deplete L22 or L22L1 for 48 h. **C.-D.** HEK293T cells were transfected with plasmids to overexpress L22 or L22L1 for 48 h. **A., C.** RT-qPCR analyses to detect the indicated target mRNAs were performed, with normalization to *36B4*. Statistical significance: ns, not significant; *, P≤0.05; **, P≤0.01; ***, P≤0.001; ****, P≤0.0001. **B., D.** Western blot analyses of the proteins indicated were performed, using GAPDH (**B)** and β-actin (**D**) as sample controls.

**Supplementary Figure 4:**
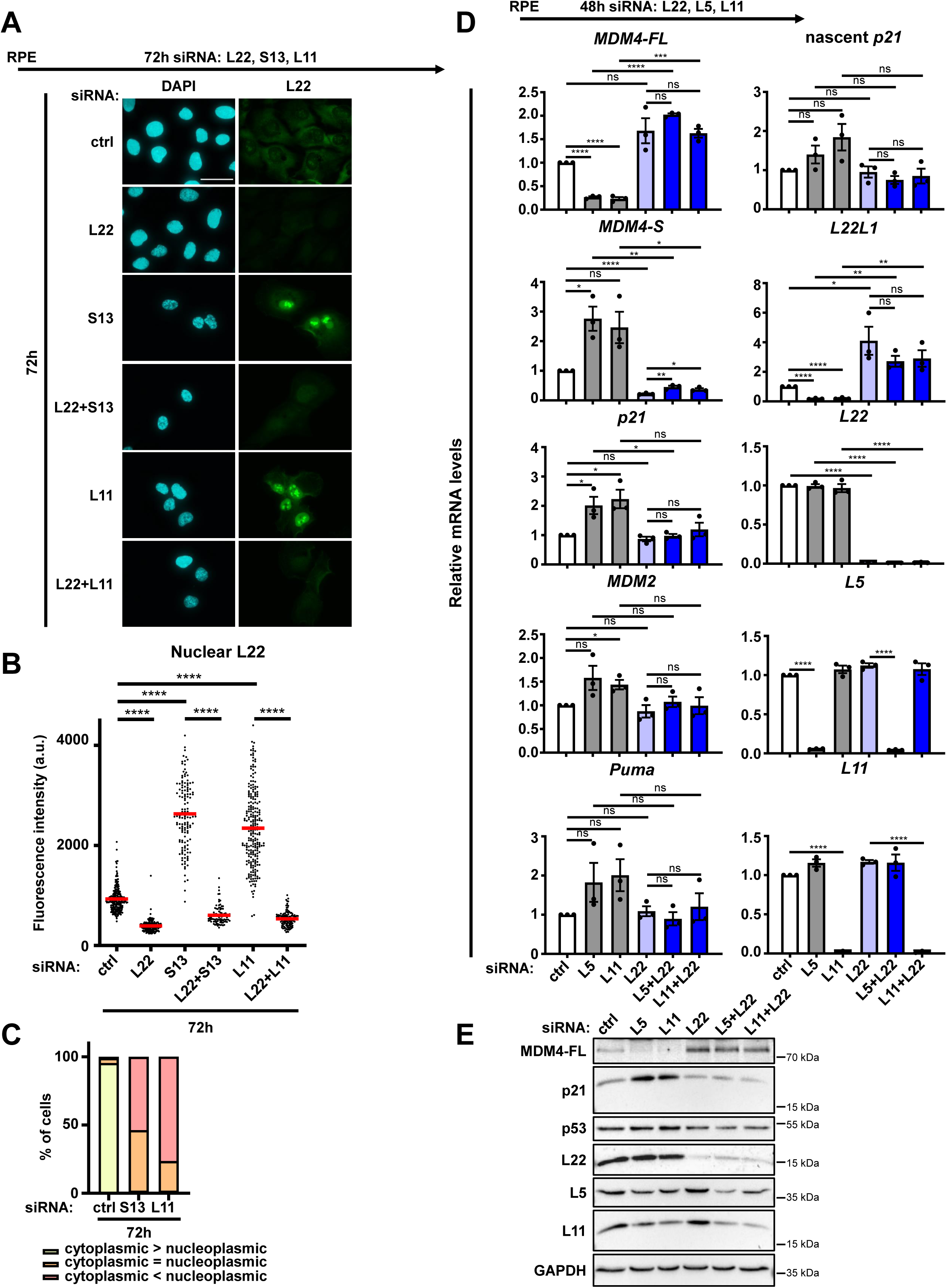
p53 activity as a function of L22 upon nucleolar stress. **A.-C.** RPE cells were transfected with siRNA to deplete L22 and/or S13 or L11 for 72 h. Immunofluorescence staining of L22 was performed. **A.** Images (100x objective, bar: 40 µm) of the nuclear (DAPI, blue) and L22 (green) staining. **B.** Quantification of single nuclear intensities of the L22 signal. Red lines represent the mean fluorescence intensity values. A minimum of 90 cells were quantified for each condition. The statistical significance was assessed by a Mann-Whitney U test: ns, not significant; *, P≤0.05; **, P≤0.01; ***, P≤0.001; ****, P≤0.0001. **C.** Quantification of cytoplasmic versus nucleoplasmic intensity of L22 signal. At least 100 cells were analyzed for each condition. **D.-E.** RPE cells were transfected with siRNA to deplete L22 and/or L5 or L11 for 48 h (the latter two to induce nucleolar stress). **D.** RT-qPCR analyses with normalization to *36B4*. **E.** Western blot analyses. Note that the depletion of L5 also diminishes the levels of L11 and vice versa, presumably due to mutual stabilization of the two proteins within the 5S RNP.

**Supplementary Figure 5:**
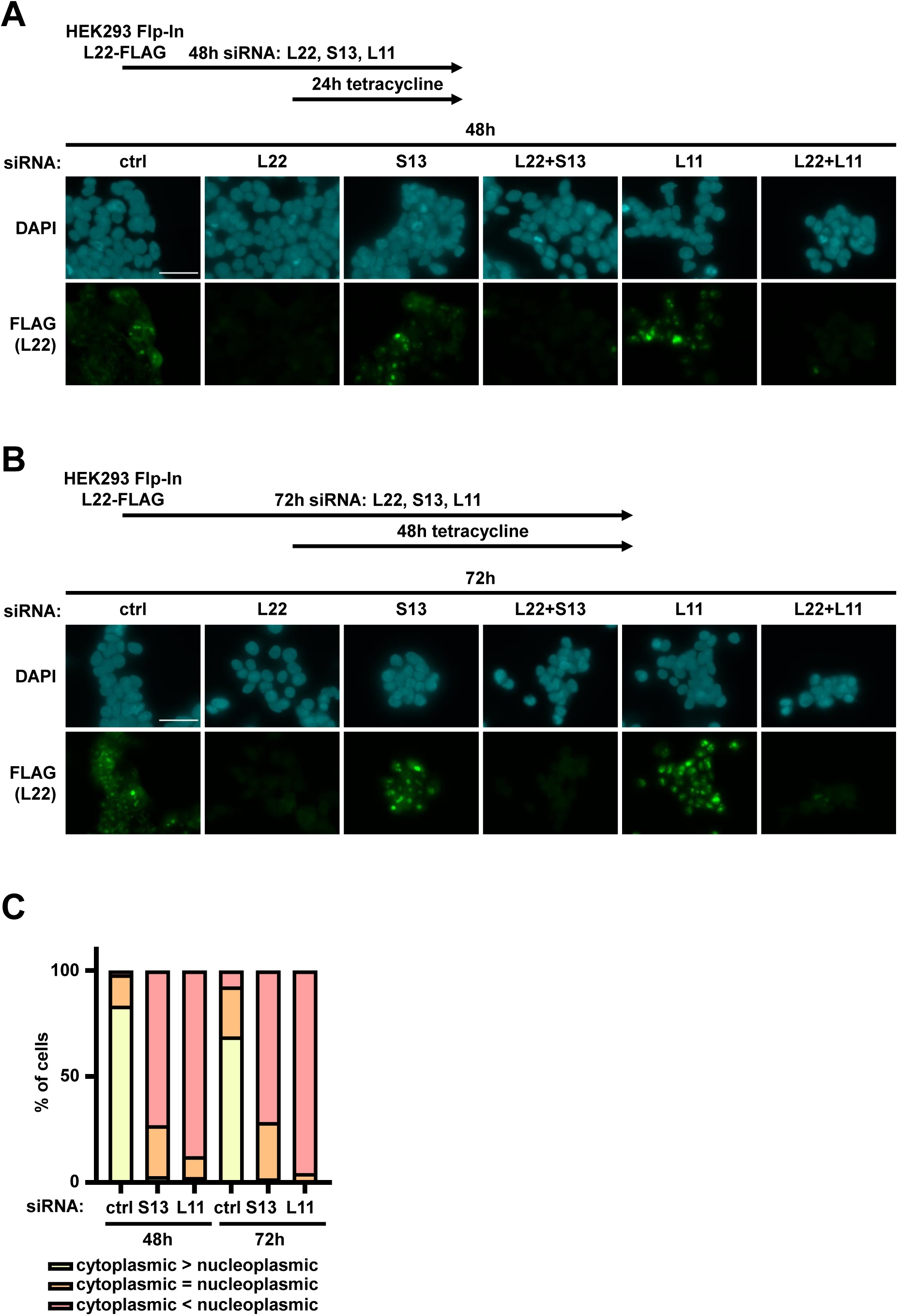
L22-FLAG localization to the nucleoplasm upon depletion of S13 or L11. **A.-C.** Stably transfected HEK293 Flp-In cells for the inducible expression of L22-FLAG were transfected with siRNA to deplete L22. To cause nucleolar stress, S13 or L11 were depleted in parallel, for 48 h or 72 h. 24 h after seeding and reverse siRNA transfection, i.e. for a total duration of 24 h respectively 48 h, tetracycline (1 µg/mL) was added for all conditions including the control, to induce the expression of FLAG-tagged L22. Immunofluorescence staining of L22 using a FLAG antibody was performed. **A.-B.** Representative images (100x objective, bar: 40 µm) of the nuclear (DAPI, blue) and L22 (green) staining upon 48 h (**A**) and 72 h knockdown (**B**). **C.** Quantification of cytoplasmic versus nucleoplasmic intensity of the L22 signal. At least 100 cells were quantified for each condition.

**Supplementary Figure 6:**
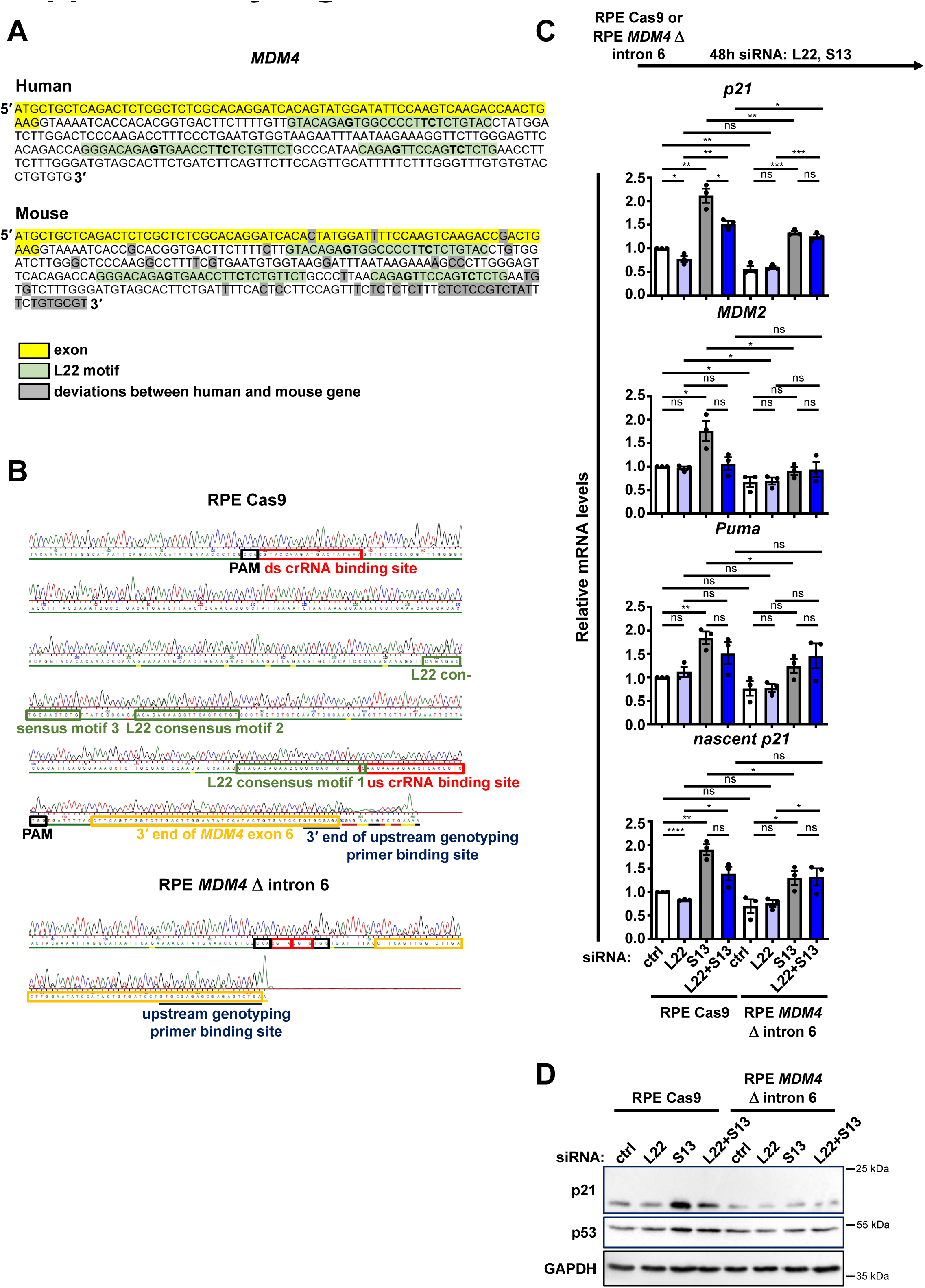
Conserved L22-binding sequences in *MDM4* intron 6 sustaining p53 activity. **A.** Sequence of exon 6 (marked in yellow) and part of intron 6 of the human and murine *MDM4* genes. Sequences in intron 6 predicted to form L22-binding stem loops are marked in green, and the characteristic bases G, C and U (T) are indicated in bold. Nucleotide differences in the murine compared to the human gene are marked in grey. **B.** Genotyping PCR products of RPE Cas9 and RPE *MDM4* Δ intron 6 cells were sequenced using a reverse sequencing primer binding at the 3′ end of the reverse genotyping primer. The following sequences are marked as also indicated in the figure: 3′ end of *MDM4* exon 6 (yellow frames), upstream (us) and downstream (ds) crRNA binding sites (RPE Cas9) respectively parts thereof (RPE *MDM4* Δ intron 6) (red frames), protospacer-adjacent motif (PAM) sequences (black frames), sequences predicted to form L22-binding stem loops (green frames). In addition, the binding site of the upstream (forward) genotyping primer respectively its 3′ end is marked with a dark blue line. **C.-D.** RPE Cas9 or RPE *MDM4* Δ intron 6 cells were transfected with siRNA to deplete L22 and/or S13 for 48 h. **C.** RT-qPCR analyses with normalization to *36B4*. Statistical significance: ns, not significant; *, P≤0.05; **, P≤0.01; ***, P≤0.001; ****, P≤0.0001. **D.** Western blot analysis.

**Supplementary Figure 7:**
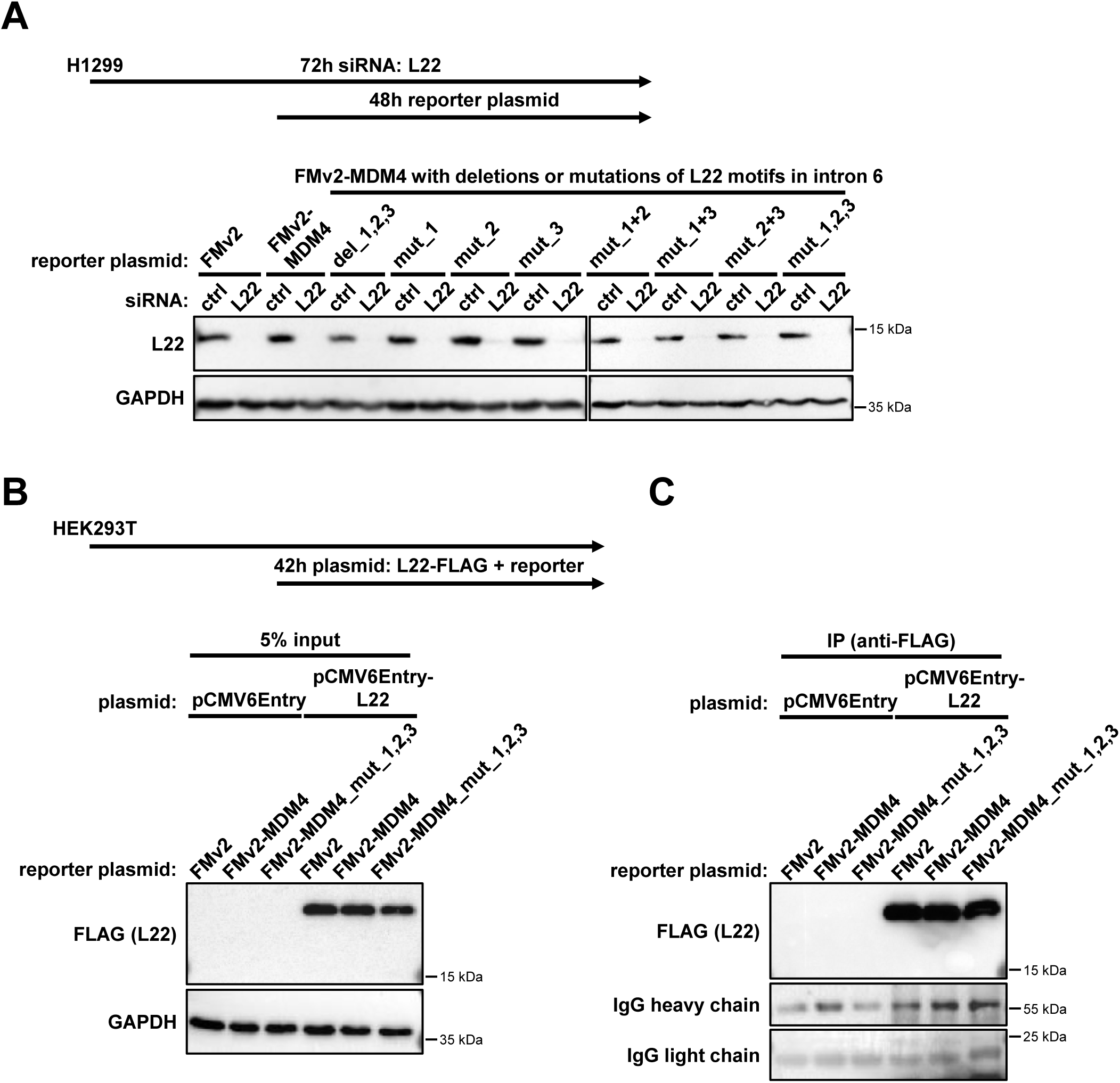
Confirmation of L22 depletion, overexpression and immunoprecipitation. **A.** H1299 cells were transfected with siRNA to deplete L22 for 72 h, and with FMv2, FMv2-MDM4 and the different mutated versions of FMv2-MDM4, at 24 hours after the siRNA transfection, followed by incubation for 48 h. Western blot analysis was performed to confirm L22 depletion, using GAPDH as a loading control. **B.-C.** HEK293T cells were transfected with pCMV6Entry or pCMV6Entry-L22 and with either FMv2, FMv2-MDM4 or FMv2-MDM4_mut_1,2,3 for 42 h, followed by RNA immunoprecipitation (RIP) using FLAG beads to precipitate tagged L22. Overexpression (**B**) and precipitation (**C**) of L22 were confirmed by Western blot analysis.

